# Detection of active Granzyme A in NK92 cells with fluorescent activity-based probe

**DOI:** 10.1101/869149

**Authors:** Sonia Kołt, Tomasz Janiszewski, Dion Kaiserman, Sylwia Modrzycka, Scott J. Snipas, Guy Salvesen, Marcin Drąg, Phillip I. Bird, Paulina Kasperkiewicz

**Affiliations:** Wrocław University of Science and Technology, Department of Chemistry, Wyb. Wyspiańskiego 29, 50-370 Wroclaw, Poland; Monash University, Monash Biomedicine Discovery Institute, Department of Biochemistry and Molecular Biology, 23 Innovation Walk, Clayton VIC 3800, Australia; NCI-designated Cancer Center, Sanford-Burnham Prebys Medical Discovery Institute, La Jolla, CA 92037, USA

## Abstract

Cytotoxic T-lymphocytes (CTLs) and natural killer cells (NKs) kill compromised cells to defend against tumor and viral infections. Both effector cell types use multiple strategies to induce target cell death including Fas/CD95 activation; and the release of perforin and a group of lymphocyte granule serine proteases called granzymes. Granzymes have relatively broad and overlapping substrate specificities and may hydrolyze a wide range of peptidic epitopes; it is therefore challenging to identify their natural and synthetic substrates and to distinguish their localization and functions. Here, we present a specific and potent substrate, an inhibitor, and an activity-based probe of Granzyme A (GrA) that can be used to follow functional GrA in cells.

## Introduction

Granule-Associated Serine Proteases (Grs) play a pivotal role in the immune system and are released by cytotoxic NKs and CTLs to eliminate abnormal cells, such as those infected with bacteria or viruses, or cancer cells^1, 2^. Grs are stored in granules within the killer cell which, on engagement with a target cell, fuse with the killer cell membrane releasing their contents into the synaptic space. Besides Grs, granules contain the pore forming cytolytic protein (perforin). Perforin polymerizes to form a pore within the target cell membrane, allowing Grs to enter the target cell cytoplasm, which leads to dramatic changes in its morphology: nuclear changes, chromatin de-condensation, membrane blebbing and, finally, cell death by lysis^3, 4^.

There are five Grs in humans, namely, Gr A, B, H, K, and M. Grs share similar structural features and use a catalytic triad of His-Asp-Ser to accomplish substrate protein cleavage. Granzyme A (GrA) is the only Gr that forms a dimer and is located at the same branch of an evolutionary tree as Granzyme K (GrK), found in the tryptase cluster at chromosome 5. These enzymes share 39.7% similarity (with 89% certainty), representing one of the most complementary pairs of Grs. Both enzymes are tryptase-like proteases and cleave after basic amino acid residues. Surprisingly, GrA and GrB share some substrates, and among them, twenty-two were cleaved in the same region but not at the same site^5^, while GrB antibodies were found to label GrA, providing strong evidence of their similarity^6^.

Grs are stored in an inactive form within granules to prevent effector (host cell) suicide, and Grs inadvertently released from granules into the host cell cytosol can be inhibited by endogenous inhibitors. The best-known examples of these endogenous inhibitors are serpins (serine protease inhibitors), which protect cells from misdirected Grs. SerpinB9^7-9^ protects against GrB activity, while no intracellular inhibitor has yet been identified for GrA^10^; however, serpinC1 prevents GrA activity in plasma^11^.

Despite intense research, the precise role of each and every Gr has not yet been determined. This is mainly due to the lack of specific tools for the in-depth investigation of Grs. Antibodies, which are commonly used, do not discriminate between an active and inactive form of the enzyme and therefore cannot be used to evaluate the control of activation and the subsequent activity of Grs. One solution to this problem involves activity-based probes (ABPs) that target only the active version of the enzyme. To date, only a few groups have tried to make chemical functional ABPs for Grs, but the major issue was poor selectivity of these probes for individual enzymes^12^. The first ABP for GrB and GrA designed using combinatorial chemistry methods was reported by the Craik group; however, the kinetic inhibition constant rates of the designed ABPs were quite low (k_obs_/I = 460 M^-1^s^-1^ for GrB and 2000 M^-1^s^-1^ for GrA), making them problematic for use in biological assays^13^.

Here, we describe a novel ABP that will allow us to distinguish GrA from other Grs and allow for GrA detection in biological samples. Our probe is cell permeable and detects GrA in the complex environment of NK92 cell lysates, it can be applied in future research on GrA in straightforward and convenient assays.

## Results

### GrA substrate specificity

GrA possesses a broad substrate specificity and can hydrolyze many physiological substrates^14^; as well as collagens and fibronectin, GrA hydrolyzes apurinic/apyrimidinic endonuclease-1 (Ape1)^15^, high mobility group protein 2 (HMG2)^16^, histone H1^17^, nuclear lamins^18^, poly ADP-ribose polymerase (PARP-1)^2^, endoplasmic reticulum-associated SET complex and three prime repair exonuclease 1 (TREX1)^19, 20^. We started our study by establishing an extensive GrA substrate specificity profile using a set of natural and unnatural amino acids. First, we synthesized a tailored library of fluorogenic substrates that contain the same P4-P2 motif, designed based on existing data, and at P1, we incorporated various amino acid residues (the overall structure of the library was Ac-Ile-Ser-Pro-P1-ACC, where ACC is 7-amino-4-carbamoylmethylcoumarin, Ac is an acetyl group, and P1 is an individual amino acid).

Next, we tested the activity of human GrA against individual P1 library components, and our results were in accordance with existing data concerning natural amino acids. We found that GrA interacts almost exclusively with positively charged amino acid residues, with the highest activity toward ones that contain a guanidine group (L-Arg, 92% and L-Phe(guan), 100%). However, our data suggested that the GrA S1 pocket is deep, since amino acids with long side chains containing basic residues (L-Lys, 21%) were accommodated much better than those with shorter side chains and the same chemical character such as L-Dap and L-Dab. Additionally, L-Abu(Bth) was recognized better than L-Ala(Bth), which is the one methylene group shorter. Interestingly, a substrate containing a 3-amino-benzoic acid residue (3-Abz, 47%) was cleaved by GrA better than the basic L-Lys residue, 4-amino-benzoic acid residue (4-Abz, 17.2%) or 2-amino-benzoic acid residue (2-Abz, 1.8%). In 2010, the Gevaert group, using N-terminal peptide-centric proteomic technology, characterized the substrate specificity of GrA^5^. Their results demonstrated that GrA almost exclusively hydrolyzes substrates after basic residues at the P1 position (arginine and lysine), confirming previous findings. Overall, the GrA molecule is positively charged with a pI of > 9, but the negative charge of Asp189 in the S1 pocket forces the binding of positively charged residues. The S1 pocket is deep and wide and is formed by Asp189-Ser195, Gly224-Val227, Lys219-Gly221, and Ser214-Leu217^21^, reflecting its catalytic preferences for long, large and basic amino acids. We found that GrA can also accommodate acidic amino acids in this pocket; however, with 5 times lower activity (L-Asp, 17.8%) **(Fig.1A, B and Supplementary Fig. S1)**.

To test GrA specificity at the P4-P2 positions, we utilized the HyCoSuL strategy and a library containing Arg at the P1 position^22^. We observed that the GrA S2 pocket can accommodate many structurally different amino acids with different chemical properties, including polar, nonpolar, basic, and acidic residues, proline derivatives and D-amino acids. A strong preference was evident for the proline derivative L-octahydroindole-2-carboxylic acid (L-Oic, 100%): substrates containing this derivative were approximately 40% more potently hydrolyzed than the other amino acids. Oic is a proline derivative equipped with an additional flexible cyclohexane residue. Cyclohexane can form few three-dimensional shapes and easily switch between conformations; therefore, we speculate that this ring, attached to the proline, a cyclic secondary amine, improves substrate accommodation at the S2 pocket. This pocket is formed by Asp95-Arg99 and Ser214-Leu217 loops^21^. Because of the guanidine group of Arg99, this pocket can bind aliphatic amino acids better than basic amino acids^21^. Previous studies reported the preferences of GrA to different amino acids, such as hydrophobic L-Phe or L-Asn (based on PS-SCL analysis)^13, 21^ at the S2 pocket, which is in agreement with our studies; however, previous studies were carried out on the library that contained only natural amino acids^23^ **(Fig. 1A, B and Supplementary Fig. S1)**.

**Figure 1.**
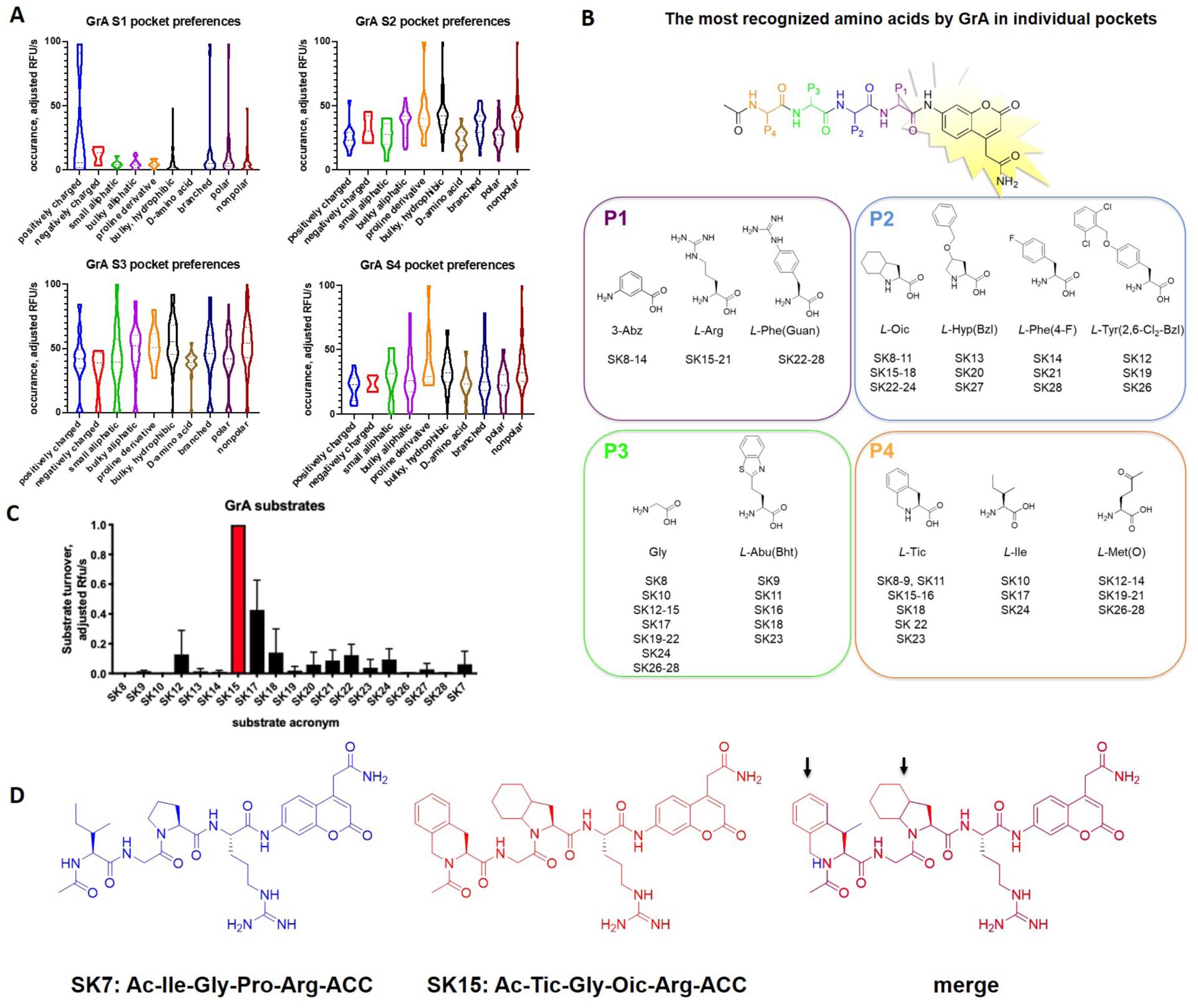
Design and characteristics of GrA substrates. **A) GrA P4-P1 catalytic preferences.** The activity of GrA was tested using the HyCoSuL strategy or individual substrates for P1 examination. Substrates were placed into the wells of the plate at the same concentration, followed by the addition of the enzyme in the reaction buffer. The increase in fluorescence was measured with 355 nm/460 nm excitation/emission wavelengths. Data are presented as a violin diagram of all events. For detailed analysis, see Supplementary Fig. 1. **B) Amino acid structures selected for new substrate synthesis based on library screening**. The best recognized sequences were chosen for individual substrate design. **C) Selection of the most active GrA substrate.** Substrates SK7-SK28 were placed into the wells of the plate at the same concentration (1 µM), followed by the addition of 89 nM enzyme in the reaction buffer. The increase in fluorescence was measured with 355 nm/460 nm excitation/emission wavelengths, accordingly. Data present the average of 2 replicates as adjusted RFU/s. **D) Comparison of SK7 (blue) and SK15 (red) structures.**

The P3 position, similar to the P4 position, has broad substrate preferences. Among a wide range of unnatural amino acids with different chemical properties, all of them, excluding D-alanine, were well recognized by GrA (>30%), whereas library substrates containing some natural residues, such as L-Arg, L-Asn, L-Glu, L-Leu, L-Lys, L-Phe, L-Thr and L-Val, were not hydrolyzed (**Fig. 1A, B and Supplementary Fig. S1**). However, the most preferred amino acid was Gly (100%), a natural, nonpolar amino acid, without a side chain. This finding is in agreement with the previous results obtained by the Gevaert group^5^; however, by extending the classic PS-SCL library with a set of unnatural amino acids, we observed that in contrast to natural sequences, all unnatural residues were recognized in this pocket. In accordance with the results of the Gevaert group, the S3 pocket of GrA is narrower than that of trypsin but is still capable of binding bulky amino acids such as L-hSer(O-Bzl) **(Fig. 1A, B and Supplementary Fig. S1)**.

The GrA S4 pocket is capable of binding various amino acids with preferences for the L-Ile from natural amino acids and for bridged analogs of L-phenylalanine, 1,2,3,4-tetrahydroisoquinoline-3-carboxylic acid (L-Tic, 100%), monooxidized methionine (L-Met(O), 77%) and threonine with a benzyl group (L-Thr(Bzl), 63%) from unnatural residues **(Fig. 1A, B)**. Methionine oxidation is a naturally occurring posttranslational modification of endogenous substrates that occurs within the organism; therefore, we speculate that GrA may be involved in the hydrolysis of posttranslationally modified substrates with L-Met(O).

### GrA fluorogenic substrate design, synthesis and kinetic evaluation

Two commercially available GrA substrates exist: Z-GPR-AFC/MNA (where **AFC** is 7-amido-4-trifluoromethylcoumarin trifluoroacetate, **MNA** is 4-methoxy-2-naphthylamide trifluoroacetate, and **Z** is a Cbz-protecting group). These substrates are tripeptides and are characterized by low kinetic constants. We therefore decided to change the peptidic leading sequence in GrA substrates. To change the sequence, based on the results from HyCoSuL library screening, we designed and synthesized eighteen fluorogenic tetrapeptides that contain ACC as a fluorophore and an acetylated *N*-terminus (named SK8-SK28), and one reference substrate based on natural amino acids (named SK7). In these sequences, we applied the combination of selected amino acids at **P4** (L-Tic, L-Ile, and L-Met(O)), **P3** (Gly and L-Abu(Bth)), **P2** (L-Oic, L-Hyp(Bzl), L-Phe(4-F), and L-Tyr(2,6-Cl_2_-Bzl)), and **P1** (L-Asp, 3-Abz, L-Arg, L-Phe(guan)) (**Fig 1B**).

We measured the activity of GrA against these new substrates and selected the top substrate, SK15 (Ac-Tic-Gly-Oic-Arg-ACC), as the leading candidate to be used in our inhibitor and ABP. GrA was approximately 40 times more active against the SK15 substrate than the reference sequence (SK7), with k_cat_/K_M_ value of 3100 M^-1^s^-1^ (**Table 1.**). Surprisingly, substrates that contain 3-Abz at the P1 position and different, defined amino acid residues at P2, P3 and P4 were not recognized by GrA, although library screening revealed that this amino acid was only two times less active than arginine. Our kinetic analyses has shown enzyme pockets cooperativity; that the binding of a particular substrate residues at a GrA S1 pocket can influence on the binding of particular residues at other subsites^24^.

**Table 1.** Steady-state Michaelis-Menten parameters for GrA, GrK and GrB determined for SK15 and SK7 substrates. *ND – no activity detected.

|  | GrA |  | GrB | GrK |
| --- | --- | --- | --- | --- |
|  | SK15 | SK7 | SK15 | SK15 |
| $K_M$ [μM] | $87.4 \pm 1.3$ | $450 \pm 48$ | ND* | ND* |
| $k_{cat}$ [$s^{-1}$] | $0.27 \pm 0.0029$ | $0.035 \pm 0.00076$ | ND* | ND* |
| $k_{cat}/K_M$ [$s^{-1}M^{-1}$] | $3100 \pm 80$ | $78 \pm 6.60$ | ND* | ND* |

Among the substrates designed by our group, sequences with L-Arg at P1 (substrates SK15-SK21) possessed the highest activity, and substrates containing L-Phe(guan) at P1 (SK22-SK28) were less active than P1-Arg substrates and more active than P1-3-Abz sequences **(Fig. 1B)**. The exchange of only one amino acid at P4 (L-Tic to L-Ile) resulted in a decrease of the hydrolysis rate (3.5 times), while the exchange of one amino acid at P3 (Gly to L-Abu(Bth)) resulted in a marked decrease in activity (63 times), suggesting that this pocket has a crucial role in substrate interactions and that the amino acid that occupies S3 should be small and aliphatic. SK15 was the most active substrate hydrolyzed by GrA, as reflected by low K_M_ values (87.4 ± 1.3 µM) and high k_cat_ rates (0.27 ± 0.0029 s^-1^) **(Fig. 1C, Table 1)**.

### The selectivity of GrA substrates (SK8-28)

Since GrA and GrK are the most structurally similar Grs, they may recognize and hydrolyze the same natural and synthetic substrates. Therefore, we tested whether our new GrA substrates are also recognized by GrK or GrB (**Fig 2A**). For this experiment, we screened substrates SK8 to SK28 and a GrK_ref_ substrate (Ac-Tyr-Arg-Phe-Lys-ACC) or GrB_ref_ substrate (Ac-Ile-Glu-Pro-Asp-ACC) at the same concentration using a plate reader, and we monitored the increase in fluorescence over time. The linear portions of the diagram were analyzed, and as we expected, the majority of the substrates were well recognized by GrK, even better than their recognition by GrA. However, the most potent substrate for GrA, SK15, turned out to be GrA selective. Additionally, we tested substrates SK8 to SK28 with GrB and noticed that GrB significantly hydrolyzed only two GrA substrates (SK13 and SK17) but was inactive on the SK15 substrate. Therefore, we concluded that Tic-Gly-Oic-Arg is a leading sequence for a specific potent inhibitor and ABP for GrA.

**Figure 2.**
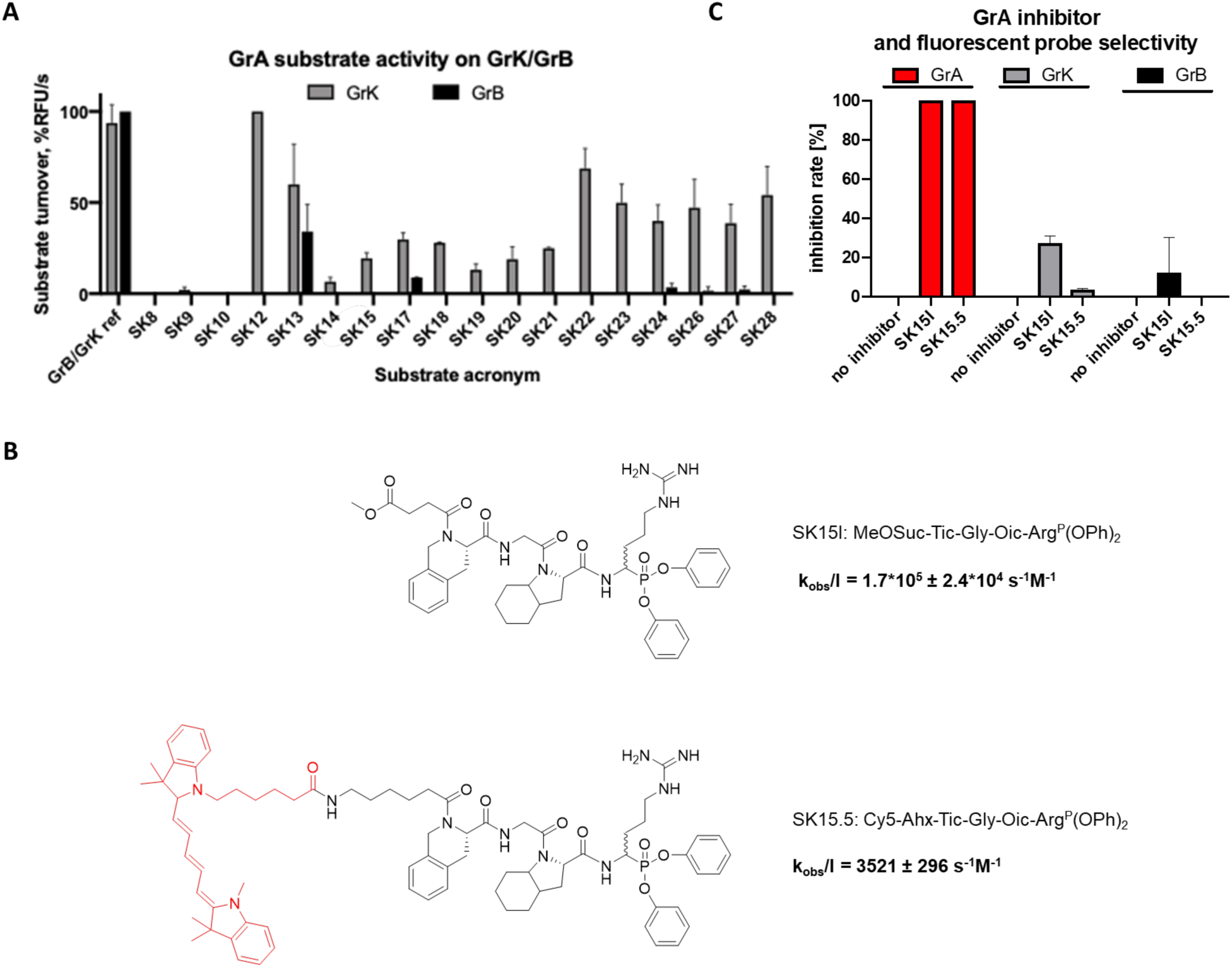
A. GrA substrate selectivity testing on GrK and GrB. Substrates SK8 - SK28, and the reference GrK and GrB substrates were placed in a 96-well plate at a concentration of 200 µM and treated with GrK or GrB. The fluorescence increase was measured with 355 nm/460 nm excitation and emission wavelengths. Data present the average of 2 replicates as adjusted RFU/s. **B. Structures of SK15I and SK15.5** and kinetic evaluation of the inhibitor and probe. **C. Selectivity of SK15I and SK15.5 toward GrK and GrB.**

### GrA inhibitor design and characteristics

Currently, biological studies on GrA activity use inhibitors characterized by low potency; or cells lacking GrA derived using gene knockout or suppression strategies. The latter strategies are time consuming and require many resources; therefore, we decided to obtain a potent and selective inhibitor that will allow us to perform so-called chemical knockdown of GrA. Based on the structure of our selective and potent substrate SK15, we designed an inhibitor with 2 main fragments: a recognition sequence and a warhead **(Fig 2B)**. Using classical techniques (solid phase peptide synthesis, peptide synthesis in solution), we synthesized an inhibitor by replacing the ACC group with an arginine derivative of diphenyl phosphonate. First, we synthesized a recognition element, based on the structure of the best substrate, by elongating the peptide on chlorotrityl resin. The *N*-terminus was protected by incorporating a methoxysuccinyl group to obtain MeOSuc-Tic-Gly-Oic-COOH **(1).** Second, we selected a diphenyl phosphonate as a reactive group that binds covalently in the active site of serine proteases. This type of inhibitor mimics the peptide structure and therefore fits perfectly within the active site of serine proteases. Using an Oleksyszyn reaction, we obtained a warhead ^+^H_3_N-Arg^P^(OPh)_2_ **(2)**. Finally, we coupled **(1)** and **(2)** in a solution, and after the coupling reaction using classic coupling reagents, we obtained MeOSuc-Tic-Gly-Oic-Arg^P^(OPh)_2_ **(3**).

We calculated the apparent second-order rate constants for inhibition (k_obs(app)_/I) of GrA-mediated SK15I under pseudo first-order conditions by serially diluting the new inhibitor in the range of 714 - 333 nM with a constant optimal substrate (SK15) concentration of 10 µM **(Fig. 2B)**. We demonstrated that the inhibitor binds with the kinetic constant of k_obs_/I = 171 000 ± 24 000 M^-1^s^-1^, which is approximately 85-fold better than the reference inhibitor (2000 M^-1^s^-1^) and is the most potent GrA inhibitor described to date.

### Fluorescent ABP design and kinetic evaluation

Antibody-related techniques allow evaluation of the total amount of enzyme present in a sample but do not distinguish between active and inactive (e.g. zymogen) forms. To date, an ABP for GrA *in-gel* detection has not been reported. This type of analysis allows detection of the active enzyme in biological samples via SDS-PAGE without the need for protein transfer and immunoblotting. We therefore designed a new probe that will allow for the convenient and straightforward *in-gel* detection of GrA in cell, tissue or fluid extracts. The probe differs from SK15I only at the *N-*terminus. The new molecule, named SK15.5 **(Fig 2B)**, was equipped with a fluorogenic moiety, the cyanine derivative Cy5, which allows the direct detection of the enzyme in samples in SDS-PAGE gels, without the need for membrane transfer. Additionally, the fluorogenic moiety was separated from the recognition sequence with an aminohexanoic acid linker, which reduces the undesirable binding of fluorophore in the active site pockets of GrA. For this purpose, first, on the chlorotrityl resin, we synthesized Cy5-Ahx-Tic-Gly-Oic-COOH and purified it with HPLC. After purification, Cy5-Ahx-Tic-Gly-Oic-COOH was coupled with the warhead molecule **(2)** to obtain Cy5-Ahx-Tic-Gly-Oic-Arg^P^(OPh)_2_. The application of a cyanine derivative at the *N*-terminus dramatically reduced k_obs_/I to 3521 ± 296 M^-1^s^-1^, which may reflect the catalytic preferences of GrA in enzyme pockets more distant from the active site. However, the k_obs_/I value for SK15.5 against GrA were still very high, and this probe may be employed for detection of the active enzyme in cells, serum, supernatants or lysates. Therefore, we tested this hypothesis in our next experiments.

### Selectivity of SK15I and SK15.5

We wondered whether the selectivity of our SK15-based molecules for GrA would be maintained by substrate structure modifications, such as replacing ACC with a diphenyl phosphonate warhead and changing the *N*-terminus from acetyl to the cyanine derivative Cy5. GrK is the only enzyme from the Grs family that hydrolyzes after basic amino acids, while GrB is one of the most potent Grs; therefore, we selected these enzymes as our selectivity controls. Focusing on this, we tested the residual activity of GrA, GrK or GrB pre-inhibited with SK15I and SK15.5 (separately) using substrates dedicated to each enzyme (SK15 for GrA, GrK_ref_ for GrK and GrB_ref_ for GrB). As indicated in **Fig. 2C**, GrK and GrB were not inhibited by our new molecules dedicated to GrA, while the activity of GrA was blocked by these inhibitors.

### *In-gel* GrA detection

Next, we tested the utility of our fluorescent probe for the *in-gel* detection of active GrA in test samples. First, we labeled purified GrA with SK15.5 at equal concentrations (the enzyme to probe ratio was 1:1) for different lengths of time in the range of 0 minutes to 2 hours. We then ran a simple SDS-PAGE assay to detect labeled GrA (**Fig. 3A**). SK15.5 is a potent probe with a k_obs_/I value of 3520 ± 296 M^-1^s^-1^; therefore, we were expecting to detect the enzyme after a short incubation time with the probe. In our hands, the probe bound to the enzyme, as indicated by the fluorescent 35 kDa species, after the minimal incubation time (5 minutes). After 30 minutes incubation, we did not observe an increase in the labeled species; therefore, we concluded that the GrA was maximally labeled with the probe. Therefore, in the next experiment, we tested the detection limit i.e. the minimum concentration of SK15.5 needed for GrA labeling, and we estimated it to be approximately 50 nM. The optimal concentration was 500 nM (strong signal) (**Fig. 3B)**.

**Figure 3.**
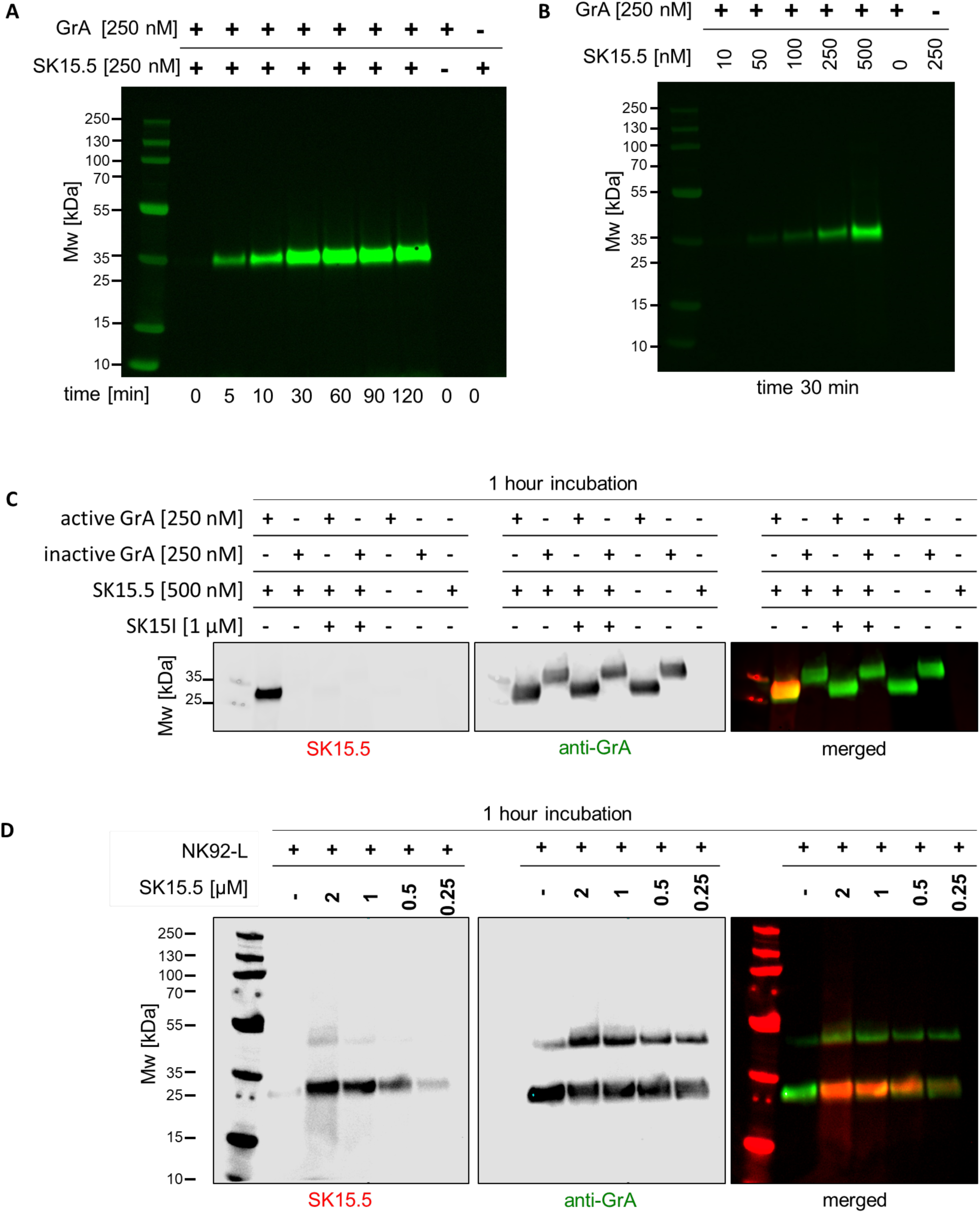
In-gel GrA detection with SK15.5 using recombinant enzyme and NK92 cell line lysates. **A) Incubation time of SK15.5 with enzyme optimization.** GrA (250 nM) was incubated with 250 nM SK15.5 for the indicated time in a range of 0-120 minutes at 37°C, followed by a reduction with 3x SDS loading buffer and SDS-PAGE analysis. **B) SK15.5 concentration optimization.** GrA (250 nM) was incubated with SK15.5 at various concentrations ranging from 10-500 nM for 30 minutes at 37°C, followed by reduction with 3x SDS loading buffer. **C) SK15.5 binds exclusively with the active GrA**. Probe binding was tested on both the active enzyme and the proenzyme at the same concentration and incubation time. The active site GrA inhibitor and the presence of prodipeptide abolished probe binding. **D) GrA in-gel detection in NK92 cell lysates.** NK92 cells in a log growth phase were lysed (1*10^*7*^ cells/mL) in lysis buffer and sonicated. Lysates were incubated with SK15.5 in a concentration range of 250 nM to 2 µM, as indicated, for 1 hour at 37°C. Samples were run on SDS-PAGE gel, transferred to a membrane, immunostained with anti-GrA, and imaged with 488 nm and 658 nm lasers. The results are representative of at least 2 replicates.

### SK15.5 binds exclusively to the active form of GrA

In contrast to the closely related GrB, GrH, GrM and GrK, GrA is a homodimer formed by a stable disulfide–linked subunits^25^. Since the subunits are oriented in opposite directions, the surrounding area may form an exosite^25^. Therefore, we aimed to verify whether the zymogen form of GrA containing the activation prodomain peptide Glu27-Lys28^26^ or the active GrA form with the processed N-terminus would bind the SK15.5 probe. After both GrA forms were incubated with SK15.5 followed by Western blot analysis, we noticed only a sharp single species corresponding to the processed form of the enzyme, and no signal was detected in the zymogen. The pre-inhibition of either GrA form with the active site-directed inhibitor SK15I also prevented probe binding **(Fig. 3C).** Therefore, we concluded that GrA must be fully processed to demonstrate catalytic activity, and that our probe binds exclusively within the GrA active site.

### GrA labeling in NK92 cell lysates

Our ultimate goal was to produce a probe that can detect not only purified, active GrA but also detects GrA in a more complex biological samples. To test whether our probe binds selectively to GrA in whole cell lysates, we selected NK92 cells, in which Grs are abundant. We lysed the cells in the exponential growth phase and to detect GrA activity, we treated lysates with SK15.5 for 1 hour. We doubled the optimal time of GrA:SK15.5 incubation to verify whether the elongation of incubation time results in non-specific binding.

We analyzed the presence of a Cy5 signal from the SK15.5 probe and observed a strong signal from a protein between 26-35 kDa, corresponding to a monomer of GrA **(Fig. 3D)**. As a control, we ran lysates without a probe, and we did not observe a signal, meaning that the observed signal was indeed from SK15.5. Additionally, an overlay of a signal from the probe and GrA antibody confirmed the binding of SK15.5 with only GrA. An additional signal between 35-55 kDa was noticed upon antibody labeling, and we speculate that it was inactive or unlabelled GrA.

We did not observe any additional bands, even at a high probe concentration (2 µM), supporting our previous finding that SK15.5 is GrA selective We concluded that our probe can be easily employed for in-gel GrA detection in cell extracts.

### GrA labeling in live NK92 cells

NK92 is an NK-like cell line that contains abundant GrA. The main goal of this research was to obtain the first ABP for GrA *in-gel* analysis. As we demonstrated in **Fig. 3**, our probe selectively binds with GrA in a complex system of cellular lysates. To test whether our probe can be used for GrA detection in living cells we first checked the cytotoxicity of the SK15.5 probe on MDA-MB-231 cells using a classic MTS assay and demonstrated that after a very long incubation time of 24 hours, our probe was cytotoxic only at a very high concentration, which was higher than the GrA amount within the cells. Therefore, we established that SK15.5 is harmless to these cells at a concentration of 1 µM **(Fig. 4A)**.

**Figure 4.**
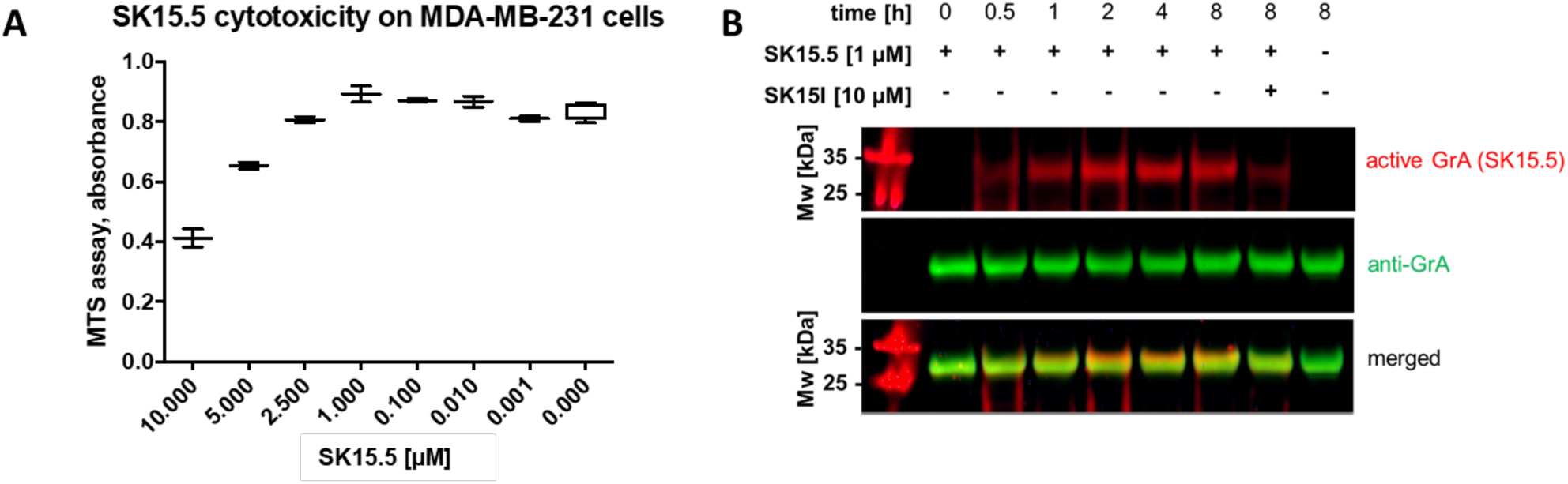
SK15.5 probe utility in living cells. **A) SK15.5 probe cytotoxicity.** MDA-MB-231 cells were treated with different SK15.5 probe concentrations in growth media for 24 hours, and the cell viability was verified with an MTS assay at 490 nm. As indicated, the probe is cytotoxic only at very high concentrations, higher than 2.5 µM. The experiment was performed in duplicate, and the data are presented as an average of these repetitions with standard deviation. **B) SK15.5 probe cell permeability.** NK92 cells in culture media were treated with SK15.5 (1 µM) for the indicated time up to 8 hours, followed by cell washing, lysis and SDS-PAGE analysis followed by protein transfer to a nitrocellulose membrane. Where indicated, the cells were preincubated with SK15I to prevent probe binding at the GrA active site. The results represent 2 biological replicates.

Next, we tested whether our SK15.5 probe is cell permeable. Thus, we treated NK92 cells with 1 µM SK15.5 for up to 8 hours and prepared cell extracts for SDS-PAGE. We noticed a strongly labeled species within 30 minutes, and the intensity increased until 2 hours **(Fig. 4B)**. This result led to the conclusion that the optimal time to detect the maximum amount of labeled GrA within NK92 cells is 2 hours, while for nonquantitative detection, 30 minutes can be used. Additionally, we utilized the GrA covalent inhibitor (10 µM, 1 hour) prior to the SK15.5 probe **(Fig. 4B, lane 7)**, and we observed a substantial decrease in probe binding. As an additional control, immunolabeling was performed and confirmed the selective binding of the probe with GrA (for full size blots, please see **Fig. S2**).

## Materials and methods

### Reagents

All chemical and biological reagents were purchased from commercial suppliers and used without further purification. The following reagents for solid phase peptide synthesis: Rink Amide RA resin (particle size 200-300 mesh, loading 0.74 mmol/g), 2-chlorotrityl chloride resin, Fmoc-protected amino acids, biotin, 1-[bis(dimethylamino)methylene]-1*H*-1,2,3-triazolo[4,5-*b*] pyridinium 3-oxid hexafluorophosphate (HATU), piperidine, diisopropylcarbodiimide (DICI) and trifluoroacetic acid (TFA) were purchased from Iris Biotech GmbH (Marktredwitz, Germany); anhydrous *N*-hydroxybenzotriazole (HOBt, purity>98%) was purchased from Creosauls; 2,4,6-collidine (2,4,6-trimethylpyridine), HPLC pure acetonitrile, and triisopropylsilane (TIPS) were purchased from Sigma Aldrich (Poznan, Poland); and *N,N*-diisopropylethylamie (DIPEA) was purchased from VWR International (Gdansk, Poland). Solvents such as N,N-dimethylformamide (DMF), dichloromethane (DCM), methanol (MeOH), diethyl ether (Et_2_O) and acetic acid (AcOH) were purchased from Avantor (Gliwice, Poland). Cy5 was purchased from Lumiprobe GmbH (Hannover, Germany). Anti-GrA antibody (rabbit, monoclonal, ab209205) was obtained from Abcam (Cambridge, Great Britain). All cell lines were purchased from DSMZ (Braunschweig, Germany) or ATCC (Manassas, USA). Peptide substrates, inhibitors and ABPs were purified by HPLC (Waters M600 solvent delivery module, Waters M2489 detector system, Waters Spherisorb S10ODS2 column, Waters sp z.o.o., Warszawa, Poland). The solvent components were as follows: water/0.1% TFA for phase A and acetonitrile/0.1% TFA for phase B. The purity of each substrate was confirmed by an analytical HPLC system using a Discovery Bio Wide Pore C8 analytical column (flow rate of 1.5 mL/min). The solvent composition was as follows: phase A (water/0.1% TFA) and phase B (acetonitrile/0.1% TFA); gradient, from 100% A to 0% A over a period of 20 minutes. The purity of all compounds was ≥95%. The molecular weight of each substrate was confirmed by high-resolution mass spectrometry on a WATERS LCT premier XE high-resolution mass spectrometer with electrospray ionization (ESI) and a time of flight (TOF) module. All kinetic assays were performed using a Spectra Max Gemini XPS spectrofluorometer (Molecular Devices) on 96-well Corning plates. The parameters of the process were as follows: 37°C; excitation/emission wavelength, 355/460 nm; cutoff, 455 nm; varying concentrations of substrates and enzyme. All measurements were performed in duplicate, and the data represent the average of these repetitions.

### Library and substrate synthesis

The P1-Arg HyCoSuL library was synthesized according to the protocols described before^27-30^. Individual substrates were synthesized using the solid phase peptide synthesis method on Rink amide resin loaded with 0.74 mmol/g. First, 3 g of resin was placed in a glass peptide synthesis vessel and swollen in DCM for 30 minutes. The *N*-terminal Fmoc-protecting group was removed with 20% piperidine in DMF solution with 5, 5 and 20 minutes cycles, and deprotection was confirmed with a ninhydrin test. Afterwards, the resin was washed thoroughly 6 times with DMF. Next, Fmoc-ACC (2.5 eq, 2.44 g) preactivated with DICI (2.5 eq, 716 µL) and HOBt (2.5 eq, 832 mg) was added to the resin, and the reaction was carried out for 24 hours. Next, the reaction was repeated with 1.5 eq of all of the above reagents to improve the yield of Fmoc-ACC coupling. Then, the Fmoc-protecting groups were removed in the same manner as above. The resin was washed 6 times with DMF, 3 times with DCM, 3 times with MeOH and dried over P_2_O_5_ overnight. Dried resin was divided into 100 mg portions and placed in a multiwell peptide synthesis vessel. To attach the P1 amino acid, 3 eq of Fmoc-P1 was preactivated with 3 eq of HATU (85 mg), and 3 eq of collidine (29 µL), and was poured onto the resin and gently agitated for 24 hours. After the deprotection cycle, the P2, P3 and P4 amino acids were coupled to the H_2_N-P1-ACC resin with DICI (2.5 eq) and HOBt (2.5 eq) as coupling reagents in coupling/deprotection cycles, preceded each time with a ninhydrin test. The *N*-terminus was protected with an acetyl group using 5 eq of AcOH, 5 eq of HBTU, and 5 eq of DIPEA in DMF. After solvent removal, the resin was washed six times with DMF to remove excess reagents, three times with DCM, and three times with MeOH, dried over P_2_O_5_ and cleaved from the resin with a mixture of TFA/TIPS/H2O (%, v/v/v, 95 : 2.5 : 2.5). The crude product was purified by HPLC and lyophilized. The purity of each substrate was confirmed by analytical HPLC and analyzed using HRMS. Substrates were dissolved in peptide-grade DMSO to a 20 mM concentration and stored at -80°C until use.

### Substrate kinetic assays

All kinetic assays were carried out in assay buffer (50 mM Tris, 100 mM NaCl, 5 mM CaCl_2_, 0.01% (v/v) Tween-20, pH 7.5). P1-Arg HyCoSuL library screening was performed on 384-well plates. Then, 0.75 µL of each library substrate was placed in one well, and preincubated (30 minutes, 37°C) enzyme solution was added. The total reaction volume was 30 µL, the final library concentration was 250 µM, and the enzyme concentration was 777 nM. Individual substrates SK8-SK28 were also tested on 384-well plates with a total substrate concentration of 67 µM and an enzyme concentration of 583 nM for GrA, 488 nM for GrK and 18 nM for GrB Briefly, substrates with an initial concentration of 20 mM were diluted 10 times in DMSO, and 1 µL of each substrate dilution was placed in separate wells on the plate, followed by 29 µL of enzyme in assay buffer. The increase in fluorescence over time was measured at excitation/emission wavelengths of 355/460 nm and a cutoff of 455 nm. The kinetic parameters of the best substrate SK15 were determined at a GrA concentration of 93 nM. Substrate concentrations ranged from 1000 µM to 7.81 µM. In the first well of the plate, 3 µL of SK15 was added to 30 µL of assay buffer, and half of this solution was transferred to a second well containing 15 µL of assay buffer. The substrate was serially diluted until eight well, in which the final substrate concentration reached 7.81 µM. Afterwards, 15 µL of preincubated (30 minutes, 37°C) enzyme was added to each well, and the fluorescence readout was measured. The K_M_, V_max_ and k_cat_/K_M_ parameters were calculated in Excel software and GraphPad Prism using Michaelis-Menten nonlinear regression.

### Inhibitor and ABP synthesis

Inhibitor and ABP synthesis was divided into two main parts. **(1)** The first step involved the synthesis of the peptide fragment (MeOSuc/Cy5-Tic-Gly-Oic-OH). 200 mg of 2-chlorotrityl resin in the peptide synthesis vessel was swollen in 10 mL of DCM for 20 minutes, and then, the resin was washed once with DCM. Next, the first amino acid, Fmoc-Oic-OH (2.5 eq, 312 mg), was dissolved in 3 mL of DCM (the mixture was cloudy) in a Falcon tube, activated for 3 minutes with 2.5 eq of DIPEA (139 µL), poured onto the resin under an argon atmosphere and stirred overnight on a platform rocker at room temperature. Then, the resin was washed 3 times with DCM, and the remaining active sites on the CTC resin were deactivated with 5 mL of methanol solution, DCM/MeOH/DIPEA (80:15:5 v/v/v), for 60 minutes. Next, the resin was washed 6 times with DMF, and the Fmoc-protecting group was removed with 20% piperidine solution in 5, 5 and 20 minutes cycles. Next, Fmoc-Gly-OH (2.5 eq, 237 mg) was coupled to the resin with HOBt (2.5 eq, 120 mg) and DICI (2.5 eq, 100 µL). Afterwards, the resin was deprotected as above, and Fmoc-Tic-OH (2.5 eq, 319 mg) was coupled in the same manner. In the case of SK15I inhibitor synthesis, the last step included the coupling of the methoxy-succinyl group (2.5 eq, 108 mg) using 2.5 eq of HOBt and 2.5 eq of DICI to protect the N-terminus of the peptide. The last step in the synthesis of the SK15.5 probe was the coupling of the Cy5 fluorophore to the N-terminus. First, the amount of chlorotrityl resin was reduced to 50 mg. Cy5 NHS ester (0.3 eq, 15 mg) was dissolved in 1 mL of DMF, preactivated with DIPEA (0.45 eq, 6.25 µL) and stirred with the resin overnight. Next, the resin was washed with DMF and dried with DCM and MeOH. The peptide was cleaved from the resin with 5 mL of the cleavage cocktail DCM:TFE:AcOH (8:1:1, v/v/v). The supernatant was collected, 20 mL of hexane was added, and the crude product was evaporated and lyophilized. The reactive warhead molecule Cbz-Arg^P^(Boc)_2_-(OPh)_2_ was synthesized according to the protocol described elsewhere^31, 32^. The Cbz-protecting group was removed using hydrogen and palladium on carbon. The obtained H_2_N-Arg^P^(Boc)_2_-(OPh)_2_ was coupled with a peptide fragment (Cy5-Ahx-Tic-Gly-Oic-COOH) with HATU (1.2 eq) and collidine (4 eq), and the reaction was monitored with analytical HPLC. When the reaction was completed, the product was extracted to ethyl acetate with 5% NaHCO_3_, 5% citric acid, and brine and dried over P_2_O_5_. Boc-protecting groups on the guanidine group of arginine were removed with a mixture of TFA:DCM:TIPS (80:15:5, v/v/v) for 30 minutes at room temperature, and TFA was removed with argon flow. The crude product was dissolved in 2 mL of DMSO, purified on semipreparative HPLC on a C8 column (Discovery Bio), lyophilized, dissolved to 10 mM in DMSO and stored at - 80°C until use.

### Inhibition kinetic assays and k_obs_/I determination

The inhibitor (SK15I) and probe (SK15.5) kinetic experiments were performed in 50 mM Tris, 100 mM NaCl, 5 mM CaCl_2_, and 0.01% (v/v) Tween-20, pH 7.5, assay buffer on a Spectra Max Gemini XPS spectrofluorometer (Molecular Devices) in 96-well plates (Corning® Costar® opaque) at 37°C. The second-order inhibition rate constant (k_obs_/I expressed in M^-1^s^-1^) toward GrA was measured under pseudo first-order kinetic conditions. Ten microliters of substrate (SK15) was mixed with varying inhibitor or probe concentrations and incubated at 37°C for 15 minutes. Afterwards, 20 µL of active GrA (350 nM) preincubated at 37°C was added, and the fluorescence increase over time was measured at the excitation/emission wavelengths of 355/460 nm and a cutoff of 455 nm. The k_obs_/I parameters were calculated in Excel software and GraphPad Prism. Experiments are presented as the average of at least three experiments with standard deviation. For residual activity assay, 488 nM of GrK or 18 nM of GrB were preincubated with 10 µM of SK15I and SK15.5 (separately) for 30 minutes at 37°C and then added to 200 µM of GrK_ref_ or GrB_ref_, respectively. As a control, enzymes without inhibitor and probe were tested. The activity was measured on spectrofluorometer with excitation/emission wavelenghts of 355/460 nm and a cutoff of 455 nm.

### *In-gel* detection of GrA using SK15.5

GrA and SK15.5 concentrations and incubation times were varied depending on the analysis performed. In the incubation time optimization assay, the enzyme and probe concentrations were constant (250 nM), and the incubation time ranged from 0 to 120 minutes. In probe concentration optimization, enzyme (250 nM) was incubated with SK15.5 at varying concentrations ranging from 10-500 nM. Incubation was carried out in assay buffer (50 mM Tris, 100 mM NaCl, 5 mM CaCl_2_, 0.01% (v/v) Tween-20, pH 7.5) at 37°C. Enzyme was incubated with probe in a volume of 80 µL, followed by the reduction with 40 µL of 3 x SDS/DTT for 5 minutes at 95°C. Samples were centrifuged (3 minutes, 4500 × g), and 40 µL of each sample was run on 4-12% Bis-Tris 10-well gels (Invitrogen). The first well was loaded with 2 µL of the protein marker PageRuler Plus Prestained Protein Ladder (Thermo Scientific). Electrophoresis was conducted at 165 V for 37 minutes at room temperature. Proteins were then transferred to a nitrocellulose membrane (0.2 µm, Bio-Rad) for 60 minutes at 10 V. Labeled proteins were detected at 658 nm (red channel) using a Sapphire Biomolecular Imager (Azure Biosystems). When testing the exclusive binding of SK15.5 with active enzyme, GrA forms (active and inactive) were diluted in assay buffer (50 mM Tris, 100 mM NaCl, 5 mM CaCl_2_, 0.01% (v/v) Tween-20, pH 7.5) to final concentration of 250 nM and preincubated with SK15I (1 µM) for 1 hour at 37°C, where indicated. Next, SK15.5 probe was added at final concentration 500 nM and incubated for another 1 hour. Total samples volume was 80 µL, to which 40 µL of 3x SDS/DTT was added and boiled for 5 minutes at 95°C. Samples were centrifuged (3 minutes, 4500 x g) and 40 µL of each sample was run on 4-12% Bis-Tris 10-well gels (Invitrogen). Each form of the enzyme without SK15.5 and probe itself were run as controls. The first well was loaded with 2 µL of protein marker PageRuler Plus Prestained Protein Ladder (Thermo Scientific). Electrophoresis and membrane transfer were conducted in the same manner as above. Membrane was blocked with 5% BSA in Tris-buffered saline with 0.1% Tween 20 (TBS-T) for 60 minutes at room temperature and immunostained overnight with rabbit anti-human monoclonal GrA antibody (Abcam, ab209205, 1:1000) and AlexaFluor 488 goat anti-rabbit secondary antibody (Invitrogen, A32731, 1:2000) for 1 hour. Membrane was imaged with 488 nm (green channel) and 658 nm (red channel) using Sapphire Biomolecular Imager (Azure Biosystems).

### GrA labeling in cell lysates

To label active GrA in the NK92 cell line (DSMZ, ACC 488), proper cell culture media had to be applied. Initially, we used three different combinations of media and supplements, according to the literature and supplier information. Cells were cultured in **(1)** alpha MEM (Sigma Aldrich), supplemented with 12.5% FBS (Sigma Aldrich), 12.5% horse serum (HS; Sigma Aldrich), 2 mM glutamine (Sigma Aldrich) and 50 U/mL IL-2 (PeproTech); **(2)** alpha MEM with 12.5% FBS, 12.5% HS, 2 mM glutamine, 0.1 mM β-mercaptoethanol (Gibco), 100 U/mL penicillin/streptomycin (Biowest) and 100 U/mL Il-2; **(3)** RPMI 1640 media (Sigma Aldrich) with 10% FBS, 2 mM glutamine, 0.1 mM β-mercaptoethanol, 100 U/mL penicillin/streptomycin and 100 U/mL Il-2. For further experiments, we chose medium **(1)** because it resulted in the highest viability of NK92 cells. Cells in the log growth phase were counted with Trypan Blue, and 10^7^ cells were collected, centrifuged, washed with DBPS (Corning®) and resuspended in 1 mL of lysis buffer (25 mM KCl, 5 mM MgCl_2_, 10 mM Tris-HCl, and 0.5% v/v Tween-20, pH 8.0). Cells were lysed by sonication (20 kHz for 10 seconds), and lysates were aliquoted and stored at -20°C until use. Samples were loaded onto 4-12% bis-Tris 10-well gels (Invitrogen), and SDS-PAGE was carried out at 165 V for 37 minutes at room temperature. Afterwards, the membrane was blocked with 5% BSA in Tris-buffered saline with 0.1% Tween 20 (TBS-T). Then, the membrane was treated with rabbit anti-human monoclonal GrA antibody (Abcam, ab209205, 1:1000) and AlexaFluor 488 goat anti-rabbit secondary antibody (Invitrogen, A32731, 1:2000) and imaged at 488 nm (green channel) and 658 nm (red channel) using a Sapphire Biomolecular Imager (Azure Biosystems).

### SK15.5 cytotoxicity assay

A cytotoxicity assay was performed with a commercially available MTS kit to determine the number of viable cells (Promega, G3580). MDA-MB-231 (ATCC®, HTB-26) cells were cultured in DMEM (Biowest) supplemented with 10% FBS and penicillin/streptomycin at a concentration of 100 IU/mL and 100 µg/mL, respectively. Cells at a density of 5×10^5^ cells/mL and a final volume of 90 µL were seeded on a clear 96-well plate 24 hours prior to the assay for proper attachment. SK15.5 probe at concentrations ranging from 1 nM to 10 µM diluted in 10 µL of DPBS was added to cells and incubated for 24 hours at 37°C with 5% CO_2_. Afterwards, 20 µL of MTS reagent was added to each well and incubated at 37°C with 5% CO_2_ for 2 hours. The absorbance was measured on a spectrophotometer (Molecular Devices) at 490 nm, and the results were analyzed with SoftMax and GraphPad Prism software.

### SK15.5 cell membrane permeability assay

NK92 cells (DSMZ, ACC 488) in the log phase were counted using Trypan Blue, and 10^6^ cells were collected in separate vials, centrifuged and resuspended in 1 mL of fresh culture media (alpha MEM supplemented with 12.5% FBS, 12.5% HS, 2 mM glutamine and 100 U/mL IL-2). To each vial containing 10^6^ cells, SK15.5 probe at a final concentration of 1 µM was added, and the vial was incubated for the indicated time (between 0 and 8 hours). As a control, GrA was preinhibited with 10 µM (final concentration) SK15I and incubated for 1 hour at 37°C, and untreated samples were simultaneously incubated. At the end of every timepoint, the cells were centrifuged, washed twice with DPBS, centrifuged and treated with 100 µL of 1x SDS/DTT solution. Afterwards, the samples were sonicated (10 kHz for 10 seconds) and loaded onto a 4-12% Bis-Tris 10-well gel (Invitrogen). SDS-PAGE was run at 165 V for 37 minutes, and membrane transfer (10 V, 60 minutes) was subsequently performed. Then, the membrane was blocked with 5% BSA in TBS-T for 60 minutes at room temperature and immunostained with rabbit anti-human monoclonal GrA antibody (Abcam, ab209205, 1:1000) and AlexaFluor 488 goat anti-rabbit secondary antibody (Invitrogen, A32731, 1:2000) and imaged at 488 nm (green channel) and 658 nm (red channel) using a Sapphire Biomolecular Imager (Azure Biosystems).

## Discussion

Grs are a family of incompletely understood serine proteases that are involved in inflammatory and apoptotic processes^1, 33^. GrA and GrK recognize the same substrate recognition motifs, and both enzymes preferentially cleave after basic amino acid residues such as arginine or lysine; however, GrA has stronger affinity for Arg, while GrK has a stronger affinity for Lys. GrA, in contrast to GrK, is a heterodimer linked by disulfide bonds, and this feature distinguishes GrA from other Grs, but has no influence on GrA/K substrate recognition motifs. These features make the design and evaluation of new chemical tools for GrA investigation challenging, and until this study, only one low potency chemical probe for GrA had been successfully designed^13^. Therefore, the goal of our studies was to develop a potent and fluorescent chemical marker for easy and straightforward labeling of GrA in its active form. Hence, we constructed the first fluorescent ABP for in-gel GrA detection and validated its utility in for labeling GrA in living cells.

GrA substrate specificity has been previously examined using PS-SCL and correlated poorly with the recognition motif in physiological GrA substrates^10, 13^. More effective searches for substrate sequence motifs are necessary; thus, in our study, we employed a HyCoSuL strategy that incorporates a broad range of unnatural amino acids to find an optimal protease substrate sequence ^22, 29, 34^.

Our findings on GrA substrate preferences correlate well with previous studies that reported natural peptidic substrates of GrA. The comparison of P1 GrA motifs determined via PS-SCL and phage display to the HyCoSuL data revealed that in addition to natural basic amino acids such as L-Arg or L-Lys, this enzyme also recognizes non-physiological residues such as phenylalanine with a guanidine group. Moreover, the enzyme recognizes these non-physiological residues five-fold better than it recognizes L-Lys, reflecting the ability of GrA to recognize bulky and positively charged amino acids. This is due to the shape and character of the S1 pocket forced by the amino acids residues forming it^21^. Despite basic amino acids, we have also observed the hydrolysis occurring after the acidic amino acid L-Asp, at a similar level as basic L-Lys; therefore, we speculate that the S1 shape of the pocket is very flexible and can adapt different amino acid residues from basic to negatively charged residues.

Previous studies reported that GrA prefers L-Asp, L-Asn, L-Phe or L-Pro P2 at the P2 position. However, when we increased the diversity of the PS-SCL library by incorporating unnatural amino acids, we noticed a dramatic increase in GrA activity on substrates with the proline derivative L-Oic. This moiety is an L-octahydroindole, and its cyclohexyl ring is flexible and tends to assume multiple nonplanar conformations. This may indicate the presence of an additional pocket close to S2 or that the shape of S2 can accommodate more complex structures than the shapes of natural amino acids. In addition to P2, our results confirm previously reported P3 and P4 specificity^5, 13^. Surprisingly, among the unnatural amino acids, incorporating bulky L-Abu(Bth) at P3 and L-Tic at P4 resulted in efficiently cleaved substrates; however, after extensive validation, substrates containing the Tic-Gly-Oic-Arg recognition motif were found to be most potently processed by GrA. We speculate that this may be due to enzyme pocket cooperativity^24^: because two neighboring residues, L-Tic and L-Abu(Bth), are bulky, there may not be enough room for both. Therefore, the application of Gly at P3 resulted in the optimal sequence design.

All synthesized GrA substrates were screened with GrK, and both enzymes hydrolyzed the majority of the substrates, confirming the similarity of the core substrate recognition motif in these enzymes. The structure of our optimal substrate (SK15) is more sophisticated and L-Oic at P2 and L-Tic at P4 allow SK15 to be well recognized by GrA and barely hydrolyzed by GrK. Therefore, we applied this recognition motif in a novel inhibitor and a fluorescent ABP, and demonstrated the first specific, potent and fluorescent marker for GrA in-gel detection.

GrA is the only member of the Gr family that occurs as a dimer. The Jenne group demonstrated the existence of the GrA dimer and the active sites orientation in opposite directions. This group also suggested the existence of exosites in the GrA homodimer near the active sites, which serve as assistants for the active site^25^. Being aware of the fact that our probe can bind to other then catalytic sites, we tested our fluorescent probe binding with active GrA and zymogen (N-terminal-processed GrA or N-terminal-containing prodipeptide GrA). We revealed that our activity-based probe did not label the zymogen form of GrA and preinhibition of active GrA with SK15I did not result in unspecific probe binding. Thus, we demonstrated that our covalent probe binds exclusively within the GrA active site.

Cells and cell lysates are more complex mixtures of proteases (mainly serine and cysteine proteases). To test the sensitivity and selectivity of SK15.5 for GrA, we incubated it with NK92 cell lysates, and we observed only one species, which was confirmed by antibody staining for GrA. We also demonstrated that SK15.5 is cell permeable and can label GrA within cells. Importantly we found that SK15.5 has very low cytotoxicity: thus, it has the potential for use in future GrA in vivo investigations.

Although Grs play a crucial role in the elimination of infected cells, their activity is also associated with disorders such as pulmonary disease, cardiovascular disease, and cancer. Furthermore, increased Grs activity correlates with patient age, since aging leads to an increase in inflammatory mediator release and to the activation of immune cells, thereby increasing Grs secretion. Therefore, in this work we have developed a novel fluorescent ABP for the fast, direct and convenient detection of GrA. In the future, this type of molecule can be used to monitor active enzyme levels within cells in targeted cell killing or cell death assays. Additionally, since SK15.5 is not cytotoxic at high concentrations and has the properties of a covalent inhibitor, it can be applied for the so-called chemical knock down of GrA (inhibition of its activity) to investigate the role and functions of GrA within cells. Finally, the fluorescent properties of SK15.5 may allow the visualization of GrA within NKs, CTLs or other cells under both physiological and pathophysiological conditions.

## Associated content

### Supporting information

The Supporting Information contains detailed screening profiles, full-size blots, amino acids structures used in libraries and individual compounds analysis for substrates, inhibitor and activity-based probe.

## Author information

### Author Contributions

S.K. and P.K. designed the project. S.K., P.K., T.J. and S.M. performed experiments. S.K. and P.K. wrote the manuscript. S.K., T.J., S.M., S.J.S., G.S., D.K. P.I.B. and P.K. reviewed the manuscript.

### Notes

The authors declare no competing financial interest.

## Acknowledgments

This work was supported by the HOMING Programme, a Grant Project of the Foundation for Polish Science, funded by the European Union under agreement No. 2016-3/24. PK is the beneficiary of L’Oreal Poland and Polish Ministry of Science and Higher Education scholarships.

## Abbreviations used

ABP: activity-based probe;
ACC: 7-amino-4-carbamoylmethylcoumarin;
AFC: 7-amido-4-trifluoromethylcoumarin trifluoroacetate;
MNA: 4-methoxy-2-naphthylamide trifluoroacetate;
HyCoSuL: hybrid combinatorial substrate library;
RFU: relative fluorescence unit;
GrA: granzyme A

