## Supplementary data for "Detection of active Granzyme A in NK92 cells with fluorescent activity-based probe"

Contents of Supporting Information

1. Purity and MS analysis of synthesized compounds…………………………………………..S2
2. Specificity profiles……………………………...…………………………………………………S24
3. Structures of amino acids…………………………………………………………….……….S25
4. Full-size blots………………………………………………………………………………...S32

**Table S1. Purity and MS analysis of synthesized compounds.**

| | Structure | $[M+H]^+$ calculated | $[M+H]^+$ measured | purity |
|---|---|---|---|---|
| **SK8** | Ac-Tic-Gly-Oic-3-Abz-ACC | 747.3142 | 747.3138 | ≥99% |
| **SK9** | Ac-Tic-Abu(Bth)-Oic-3-Abz-ACC | 908.3442 | 908.3450 | ≥98% |
| **SK10** | Ac-Ile-Gly-Oic-3-Abz-ACC | 701.3299 | 701.3295 | ≥99% |
| **SK12** | Ac-Met(O)-Gly-Tyr(2,6-$Cl_2$-Bzl)-3-Abz-ACC | 905.2139 | 905.2135 | ≥99% |
| **SK13** | Ac-Met(O)-Gly-Hyp(Bzl)-3-Abz-ACC | 787.2761 | 787.2747 | ≥99% |
| **SK14** | Ac-Met(O)-Gly-Phe(4-F)-3-Abz-ACC | 749.2405 | 749.2413 | ≥99% |
| **SK15** | Ac-Tic-Gly-Oic-Arg-ACC | 784.3782 | 784.3790 | ≥97% |
| **SK17** | Ac-Ile-Gly-Oic-Arg-ACC | 738.3939 | 738.3932 | ≥99% |
| **SK18** | Ac-Tic-Abu(Bth)-Oic-Arg-ACC | 945.4081 | 945.4095 | ≥98% |
| **SK19** | Ac-Met(O)-Gly-Tyr(2,6-$Cl_2$-Bzl)-Arg-ACC | 942.2778 | 942.2795 | ≥98% |
| **SK20** | Ac-Met(O)-Gly-Hyp(Bzl)-Arg-ACC | 824.3401 | 824.3399 | ≥99% |
| **SK21** | Ac-Met(O)-Gly-Phe(4-F)-Arg-ACC | 786.3045 | 786.3045 | ≥98% |
| **SK22** | Ac-Tic-Gly-Oic-Phe(guan)-ACC | 832.3782 | 832.3781 | ≥97% |
| **SK23** | Ac-Tic-Abu(Bth)-Oic-Phe(guan)-ACC | 993.4081 | 993.4070 | ≥97% |
| **SK24** | Ac-Ile-Gly-Oic-Phe(guan)-ACC | 786.3939 | 786.3940 | ≥99% |
| **SK26** | Ac-Met(O)-Gly-Tyr(2,6-$Cl_2$-Bzl)-Phe(guan)-ACC | 990.2778 | 990.2772 | ≥95% |
| **SK27** | Ac-Met(O)-Gly-Hyp(Bzl)-Phe(guan)-ACC | 872.3401 | 872.3403 | ≥99% |
| **SK28** | Ac-Met(O)-Gly-Phe(4-F)-Phe(guan)-ACC | 834.3045 | 834.3040 | ≥99% |
| **SK7** | Ac-Ile-Gly-Pro-Arg-ACC | 684.3469 | 684.3463 | ≥95% |
| **SK15I** | MeOSuc-Tic-Gly-Oic-$Arg^P(OPh)_2$ | 844.3799 | 844.3809 | ≥95% |
| **SK15.5** | Cy5-Ahx-Tic-Gly-Oic-$Arg^P(OPh)_2$ | 654.3615 | 654.3613 | ≥95% |

**SK8, Ac-Tic-Gly-Oic-3-Abz-ACC**

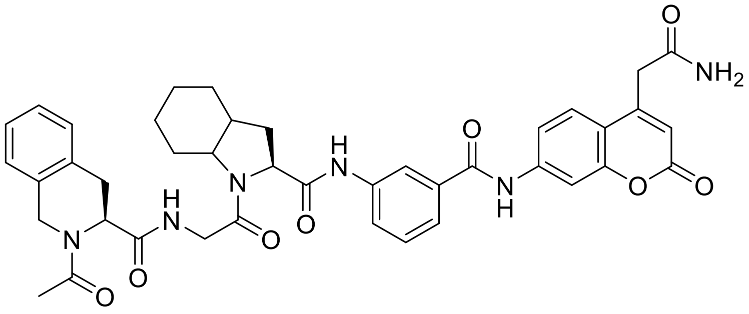

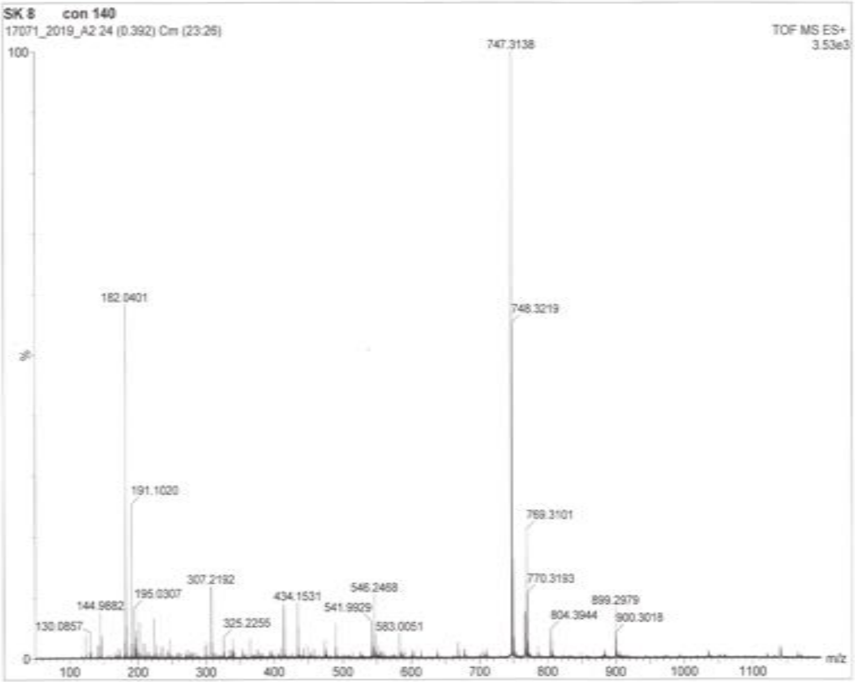


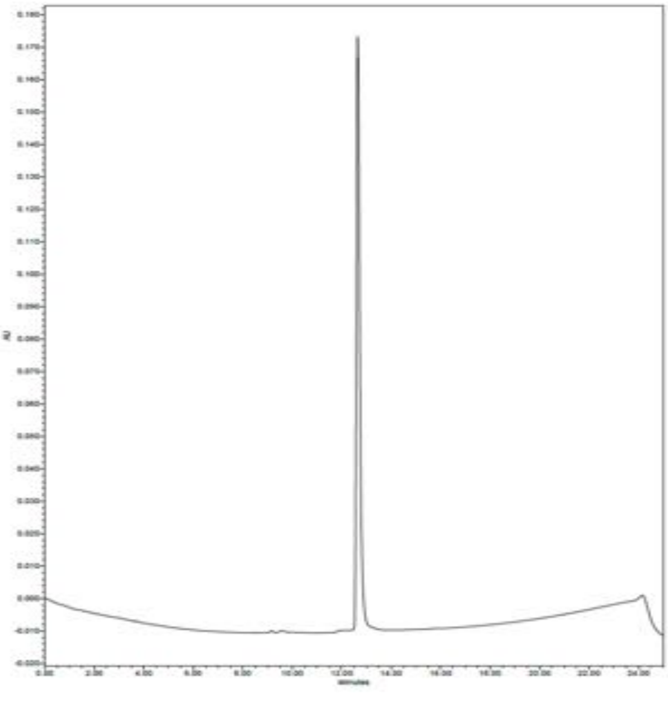

**SK9, Ac-Tic-Abu(Bth)-Oic-3-Abz-ACC**

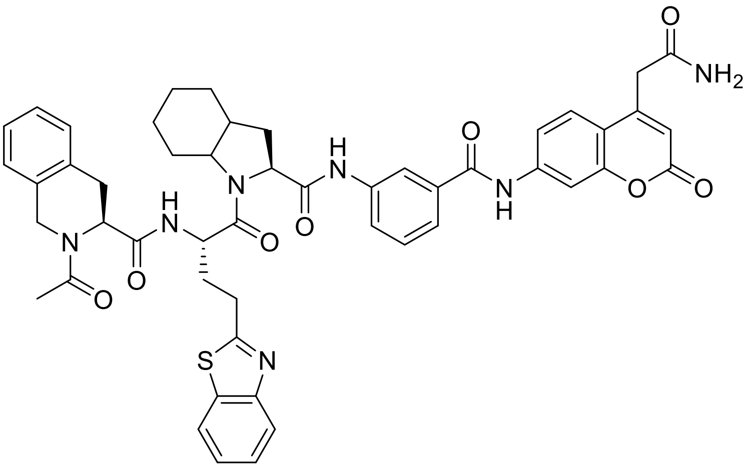


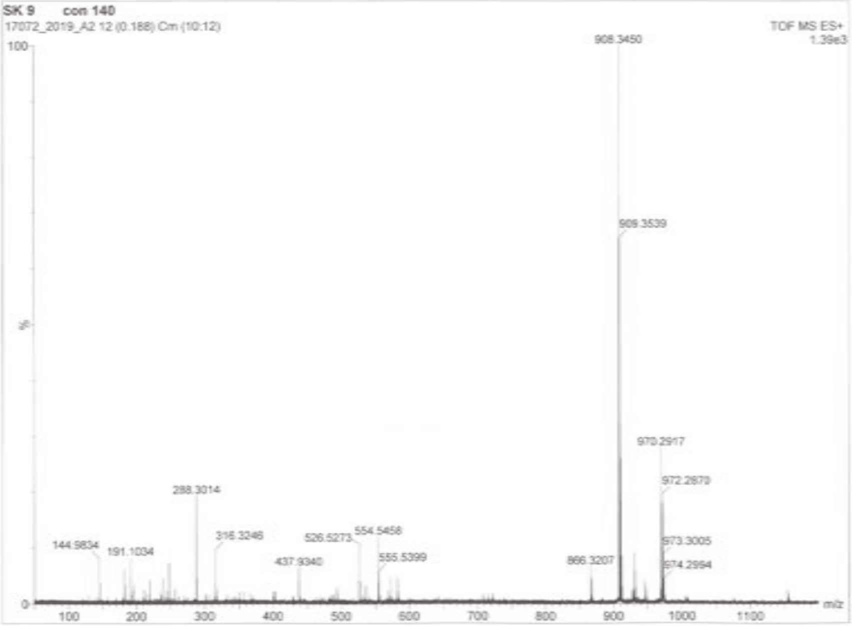


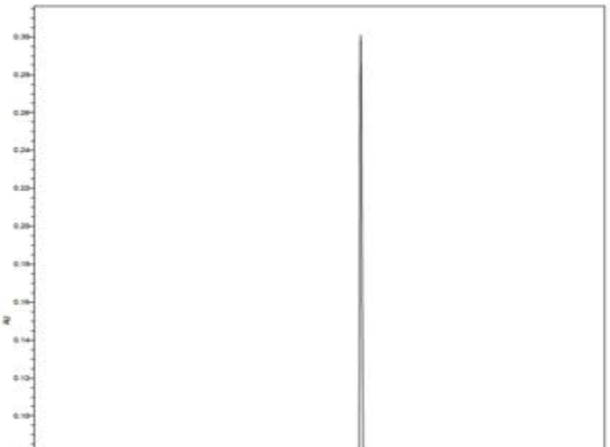

**SK10, Ac-Ile-Gly-Oic-3-Abz-ACC**

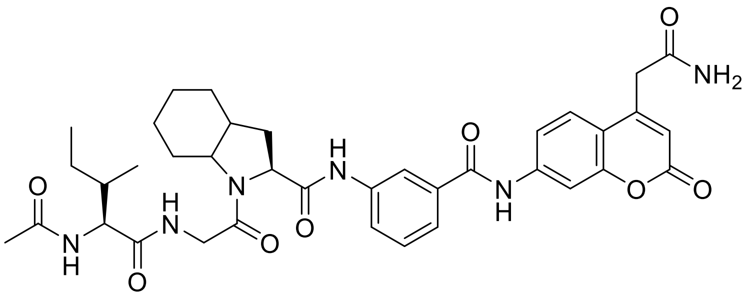

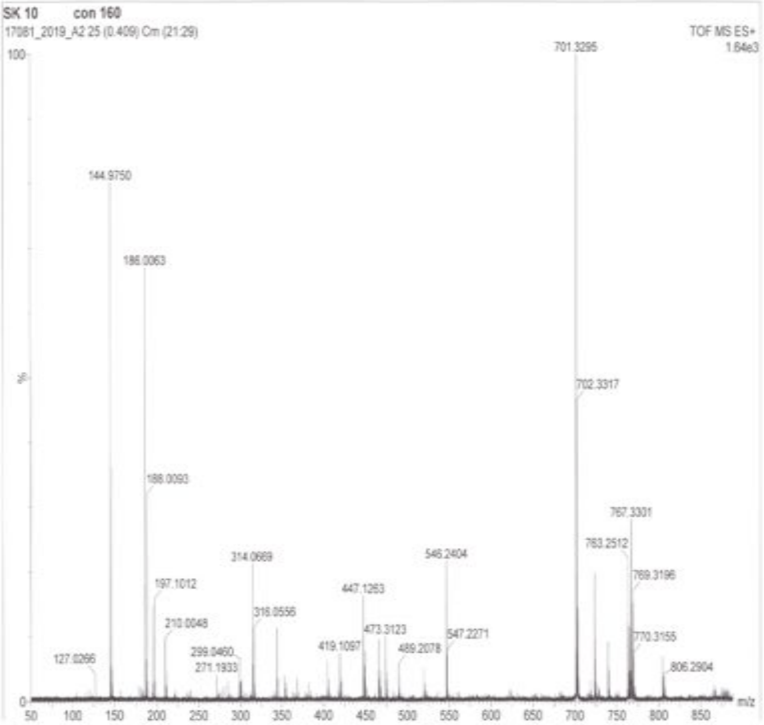


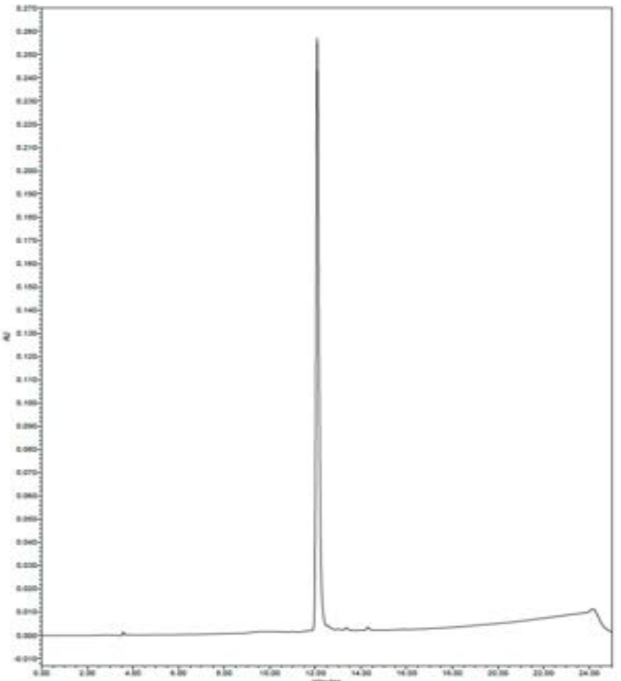

**SK12, Ac-Met(O)-Gly-Tyr(2,6-$Cl_2$-Bzl)-3-Abz-ACC**

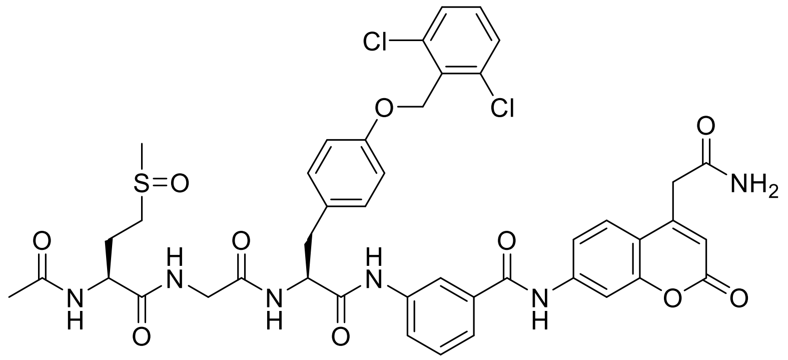

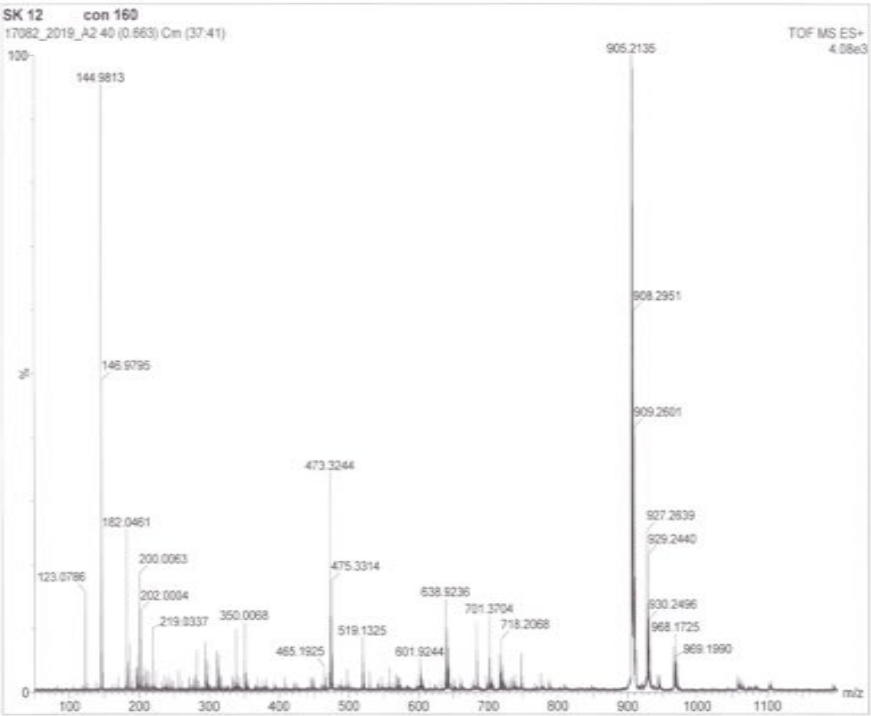


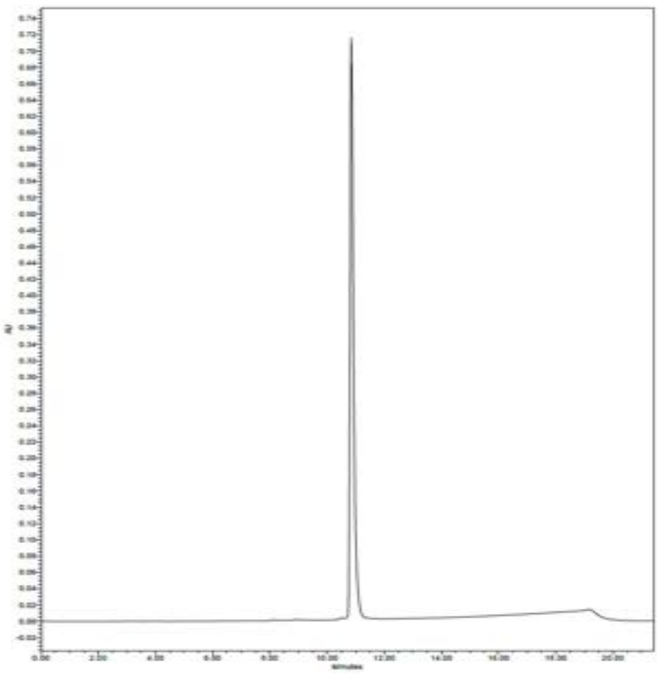

**SK13, Ac-Met(O)-Gly-Hyp(Bzl)-3-Abz-ACC**

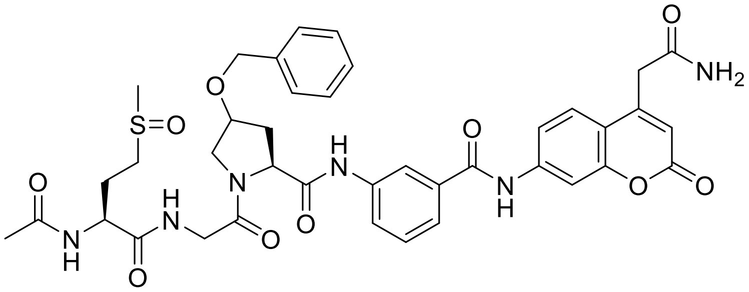

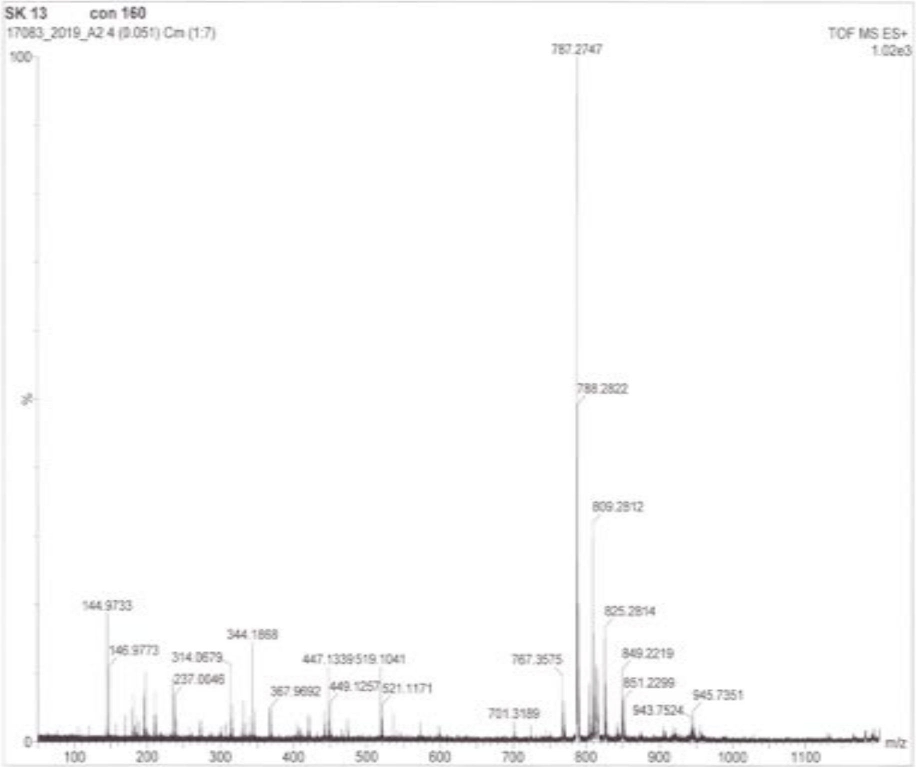


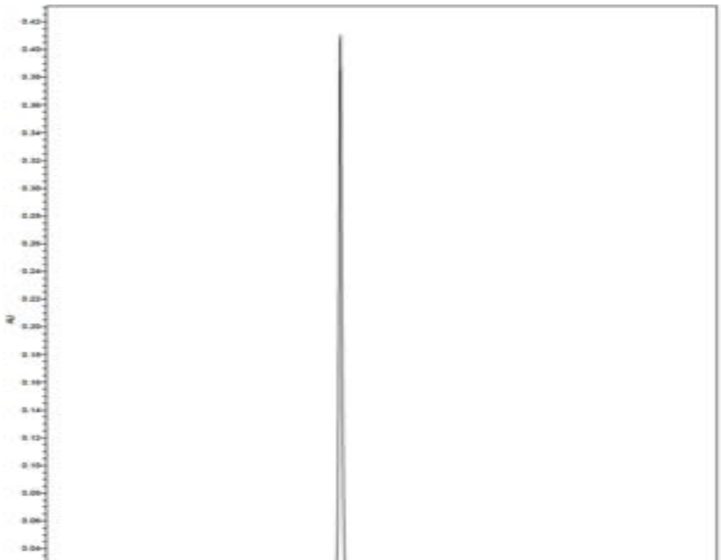

**SK14**, **Ac-Met(O)-Gly-Phe(4-F)-3-Abz-ACC**

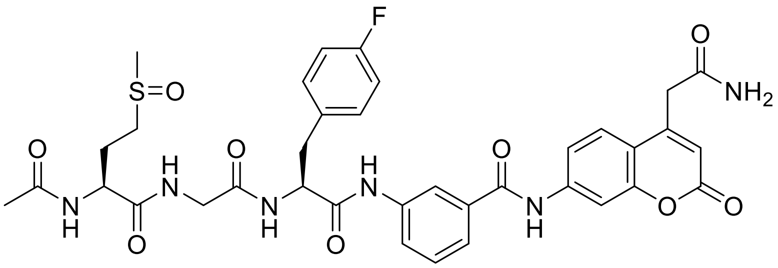

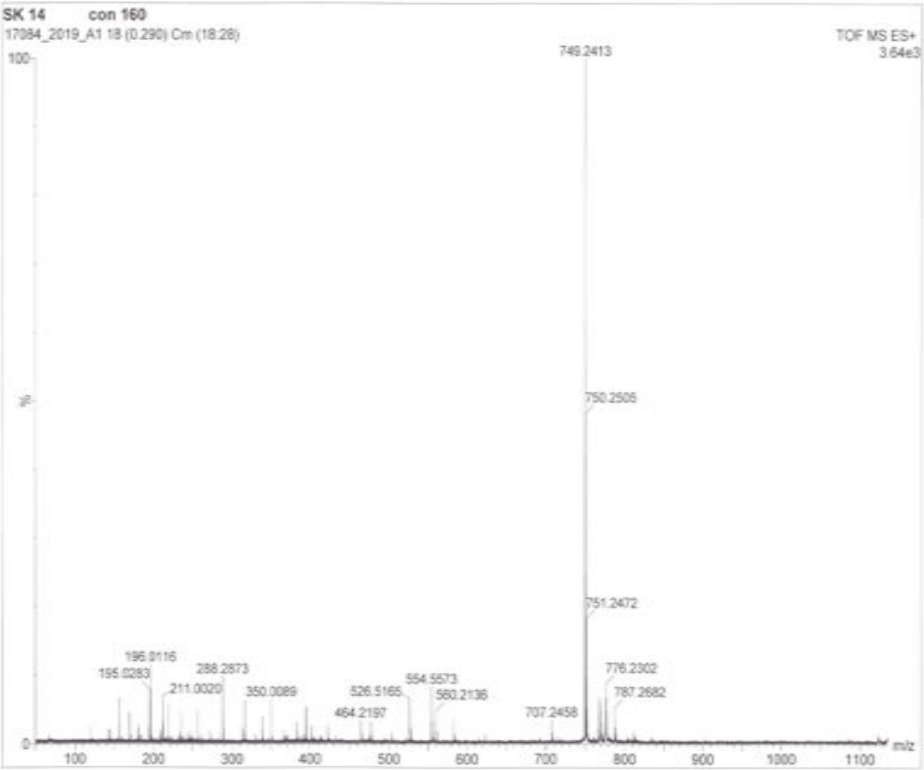


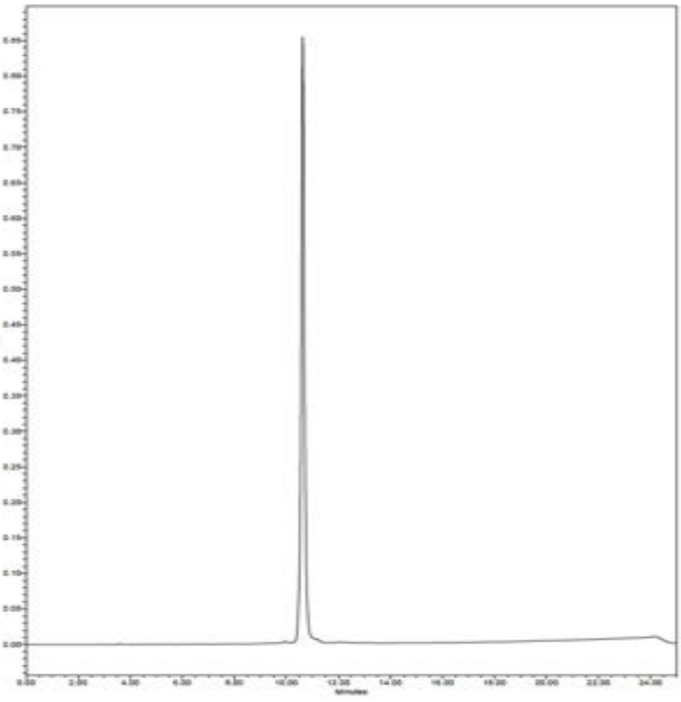

**SK15, Ac-Tic-Gly-Oic-Arg-ACC**

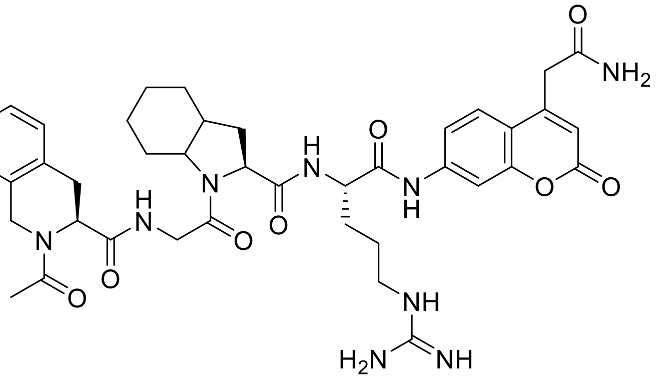

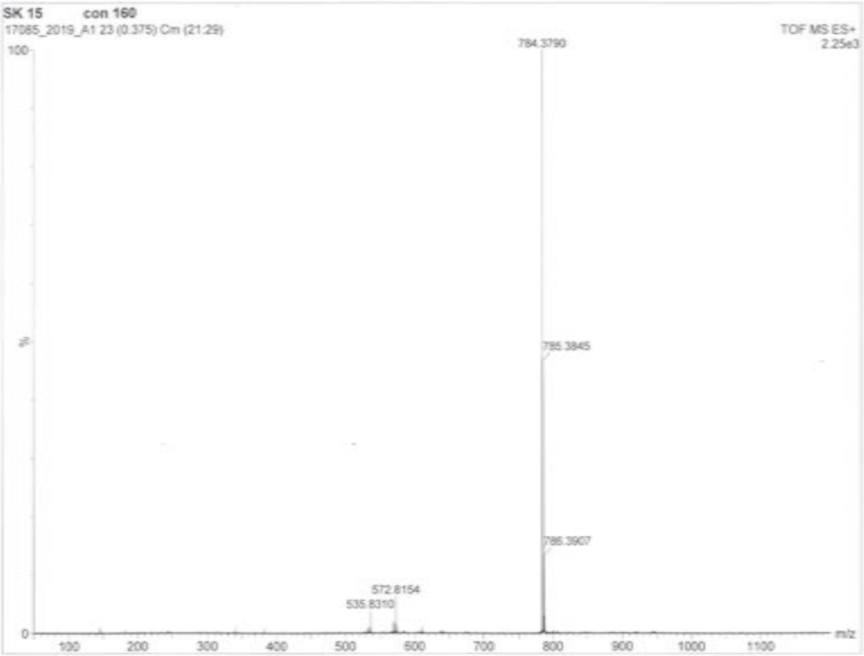


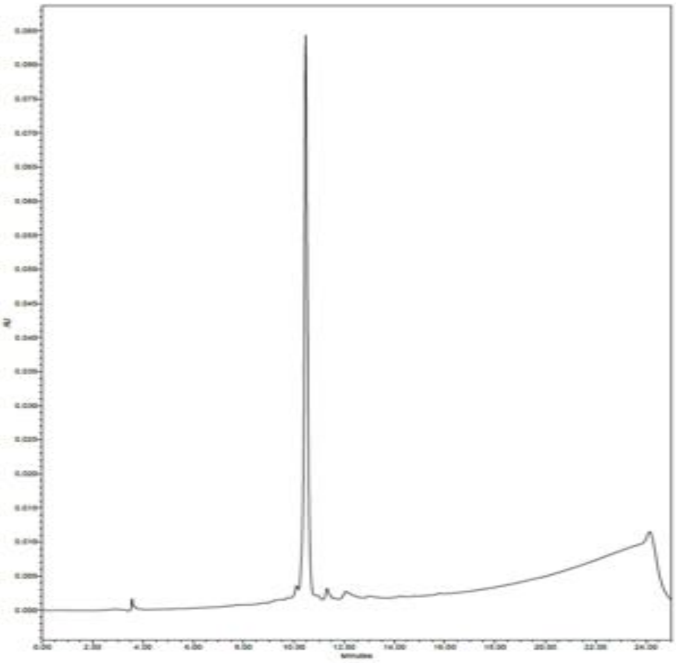

**SK17, Ac-Ile-Gly-Oic-Arg-ACC**

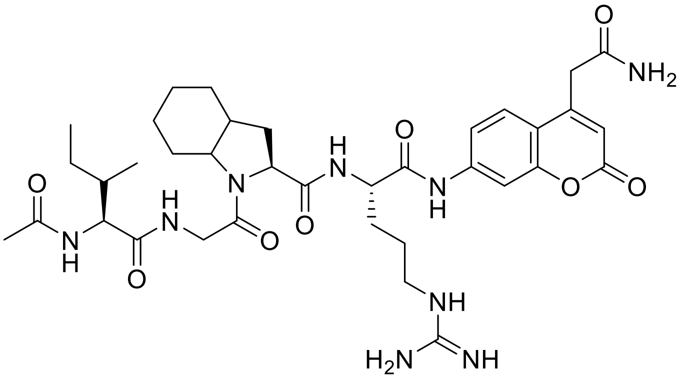


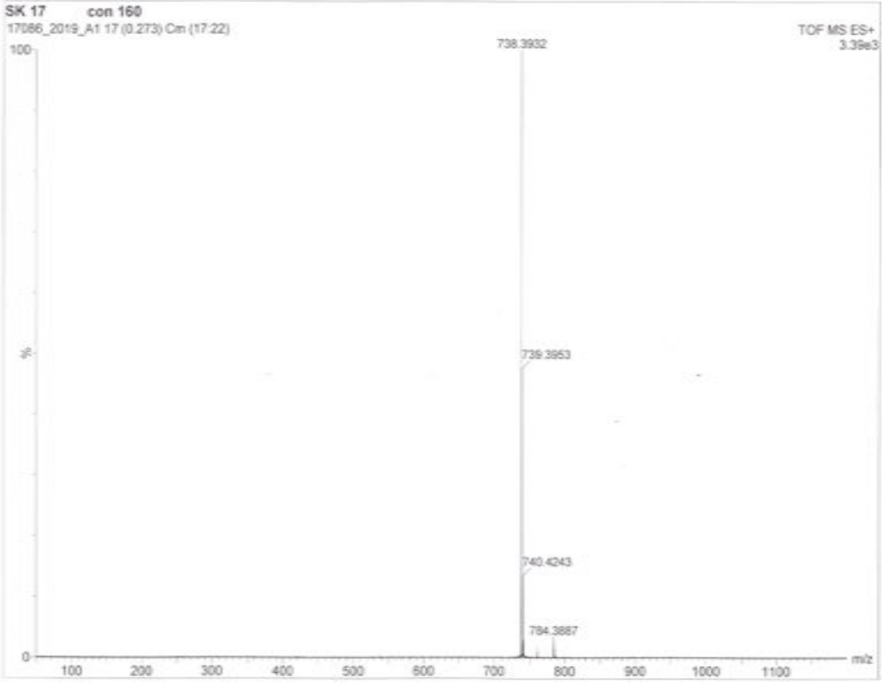


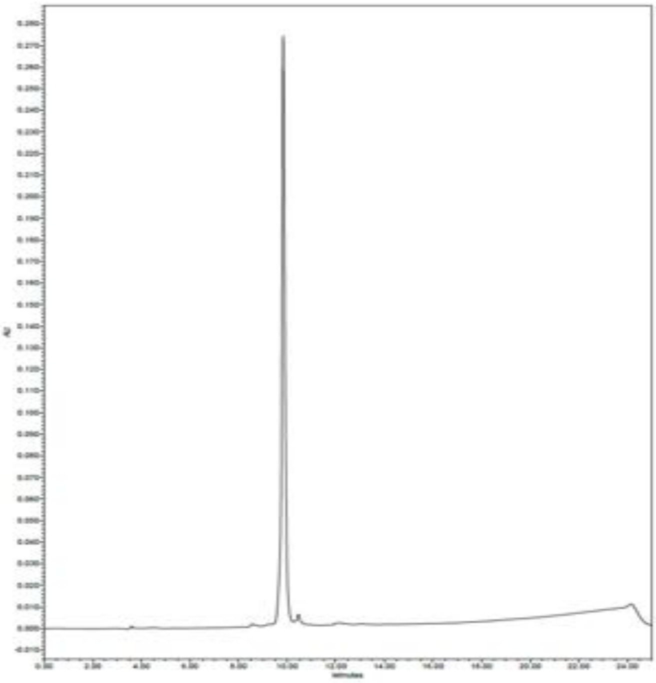

**SK18, Ac-Tic-Abu(Bth)-Oic-Arg-ACC**

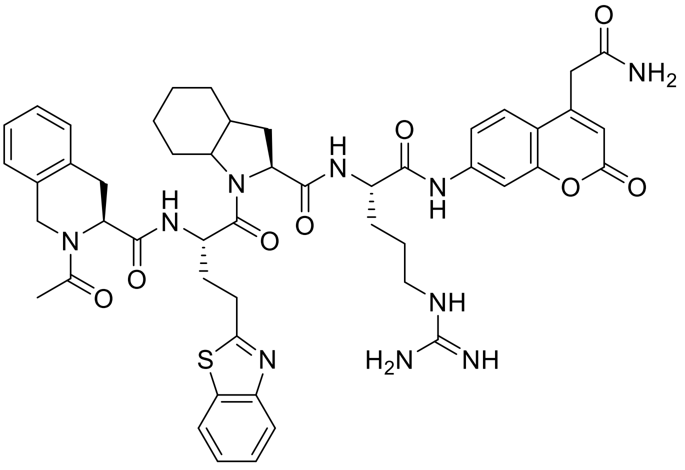

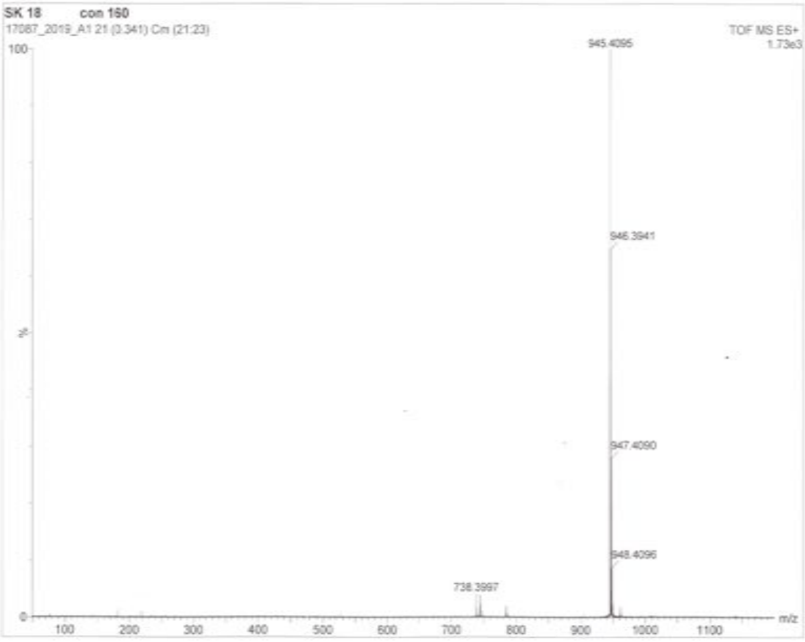


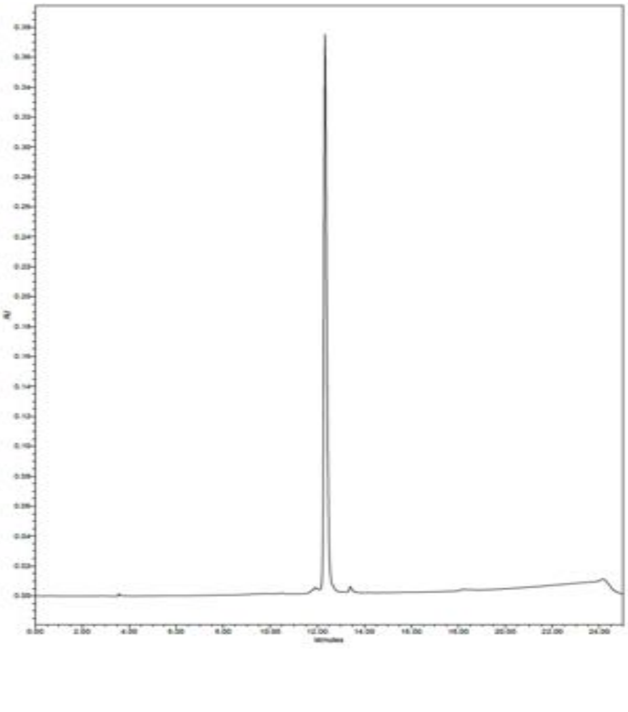

**SK19, Ac-Met(O)-Gly-Tyr(2,6-Cl$_2$-Bzl)-Arg-ACC**

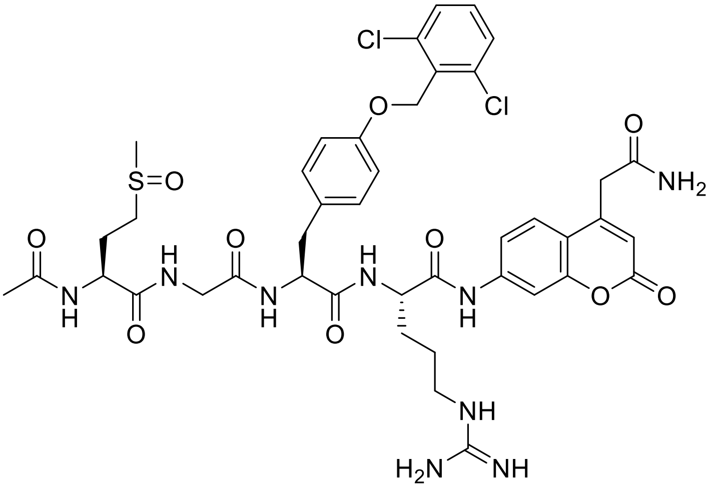


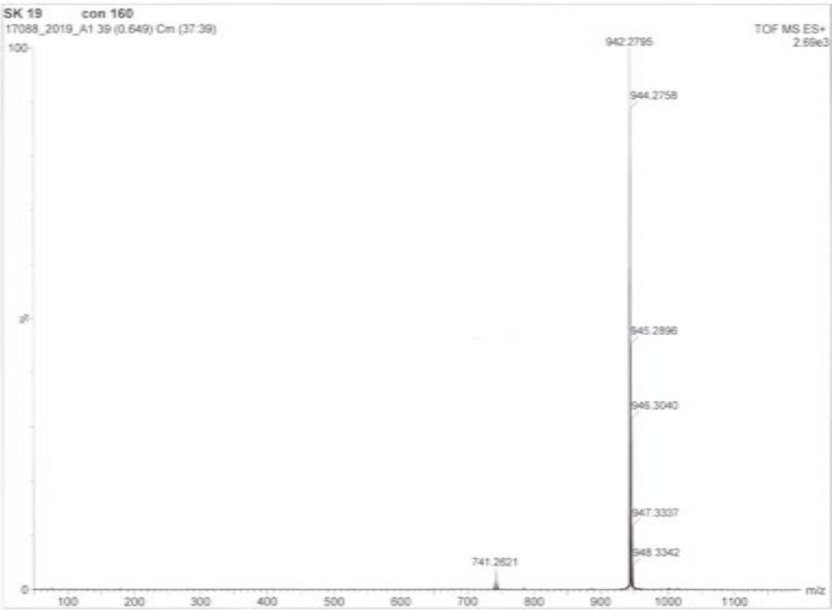


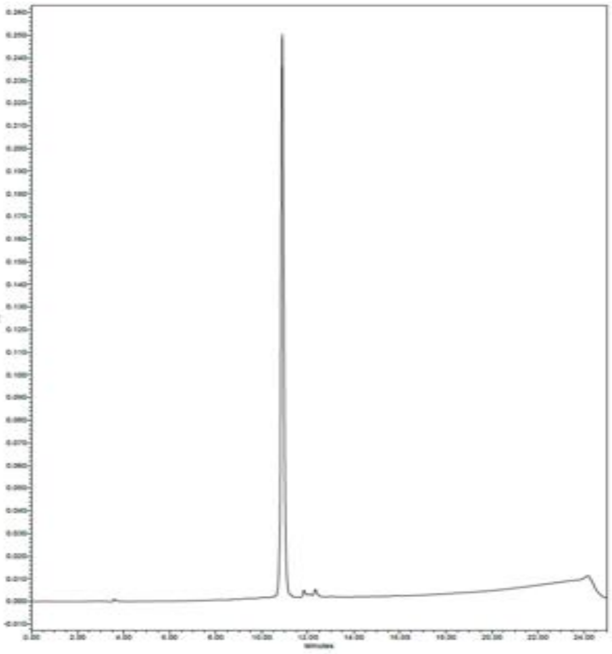

**SK20, Ac-Met(O)-Gly-Hyp(Bzl)-Arg-ACC**

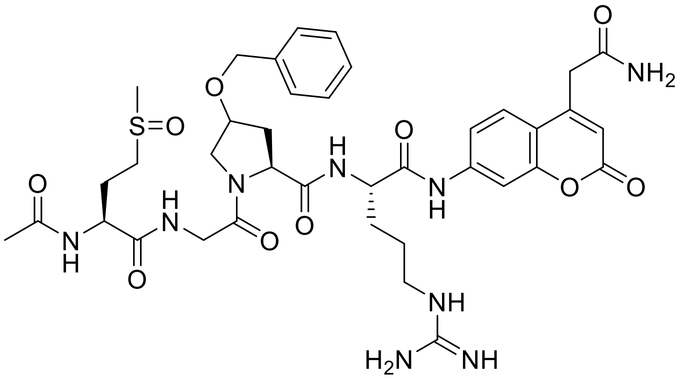

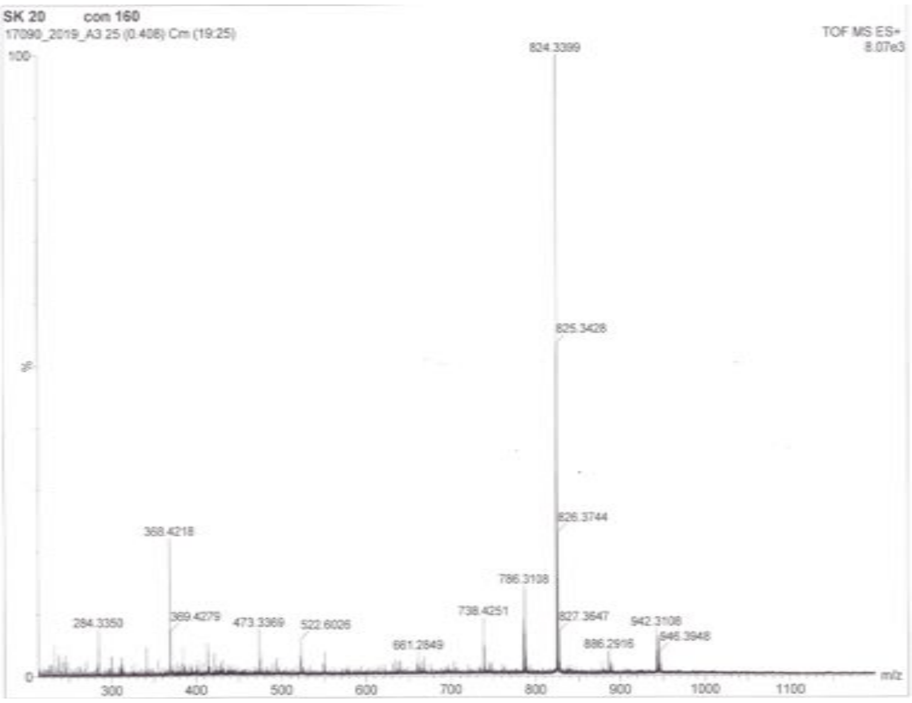


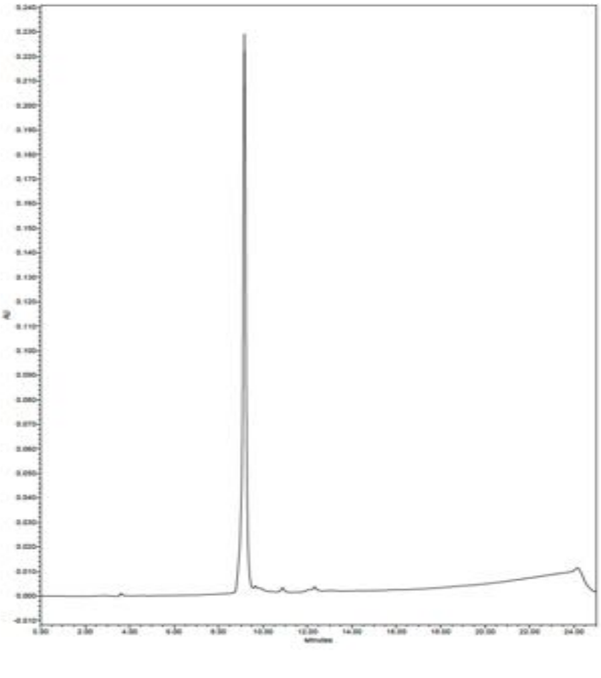

**SK21, Ac-Met(O)-Gly-Phe(4-F)-Arg-ACC**

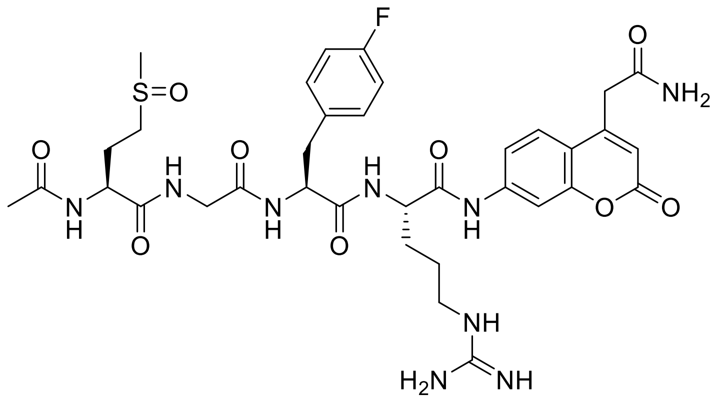

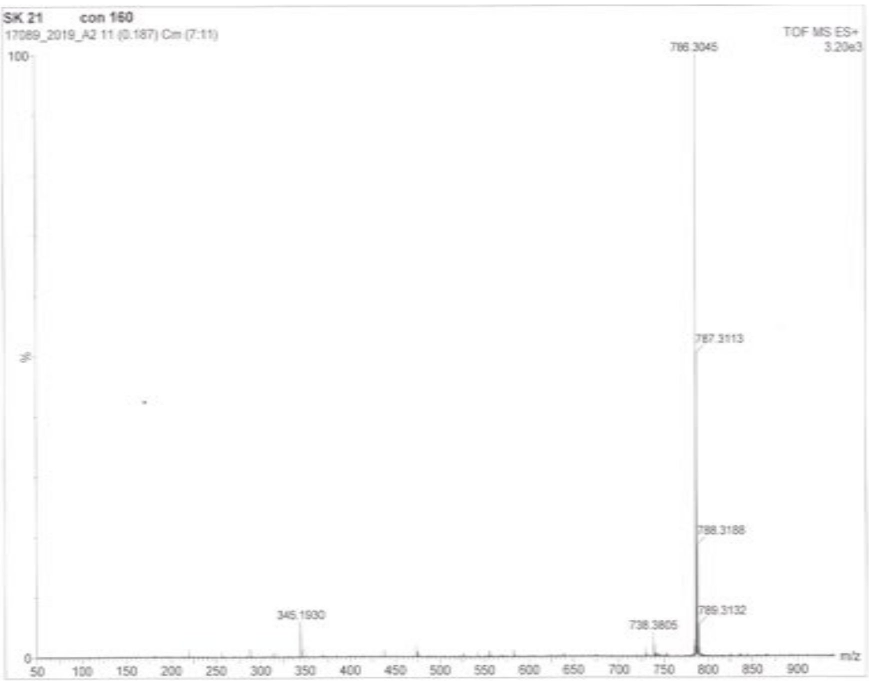


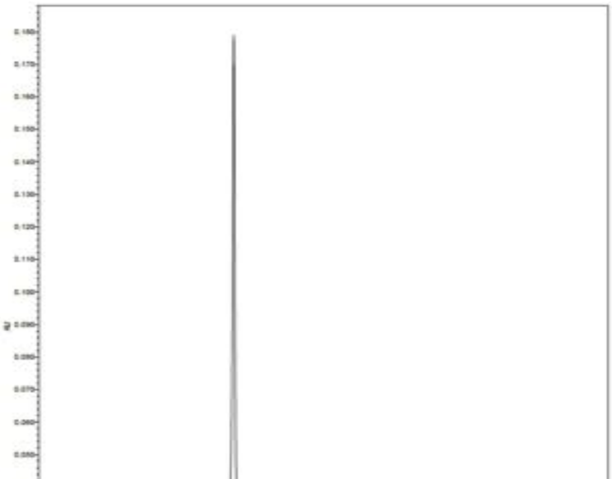

**SK22, Ac-Tic-Gly-Oic-Phe(guan)-ACC**

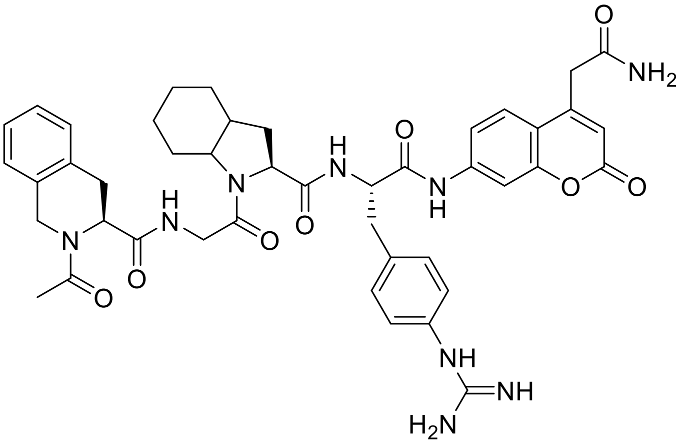

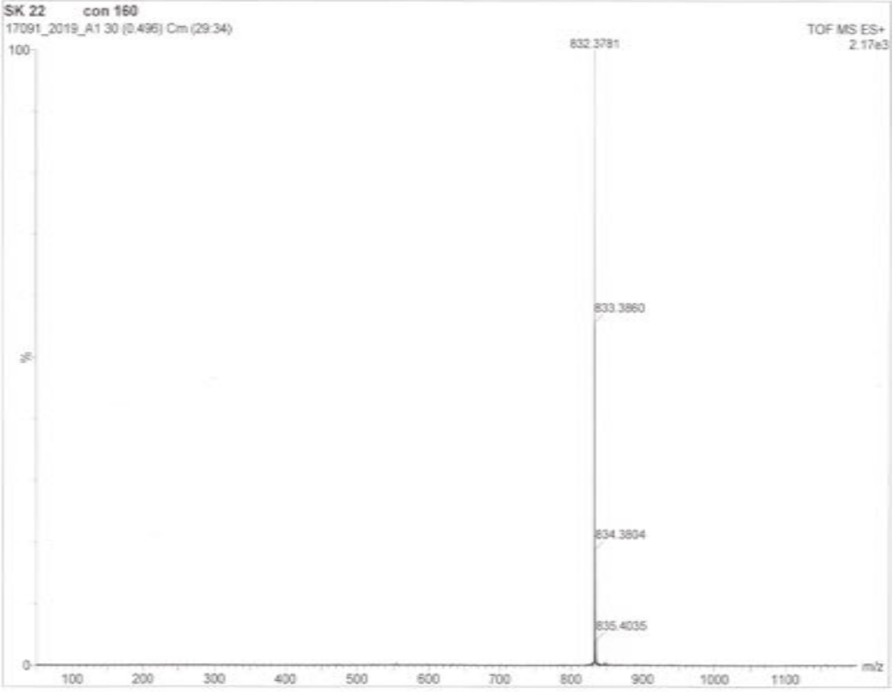


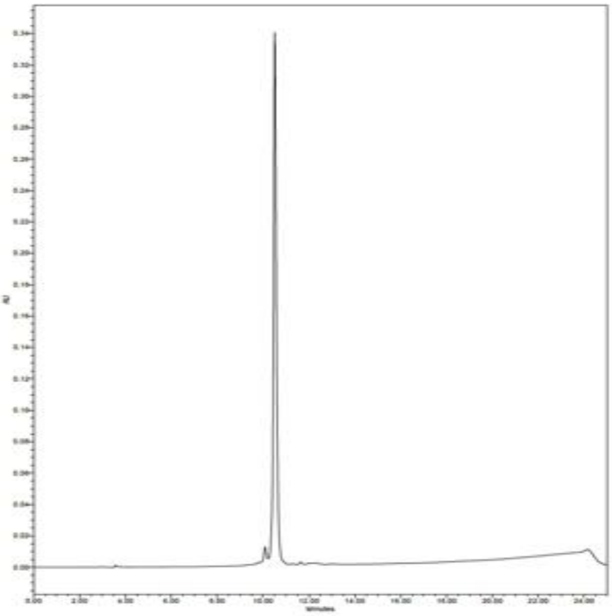

**SK23, Ac-Tic-Abu(Bth)-Oic-Phe(guan)-ACC**

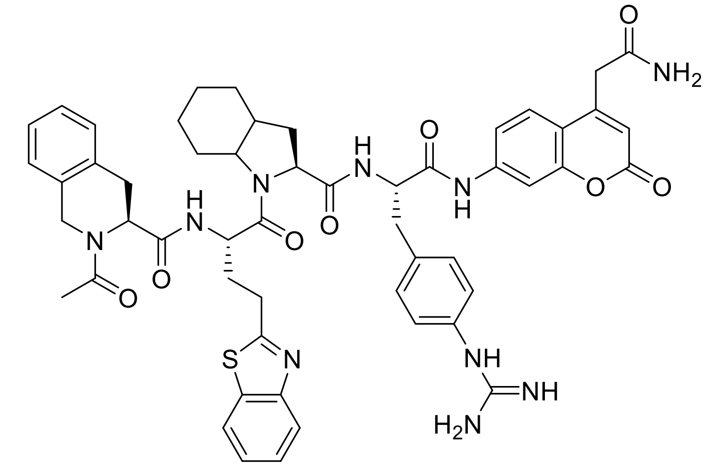


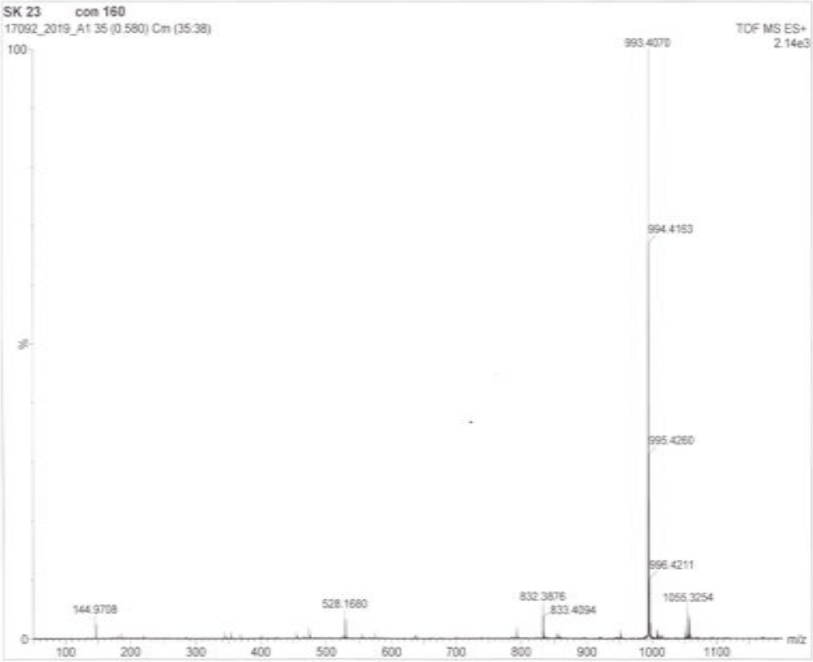


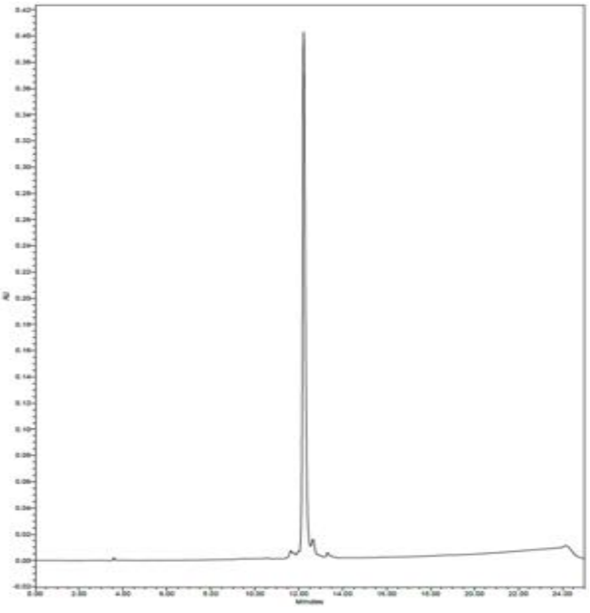

**SK24, Ac-Ile-Gly-Oic-Phe(guan)-ACC**

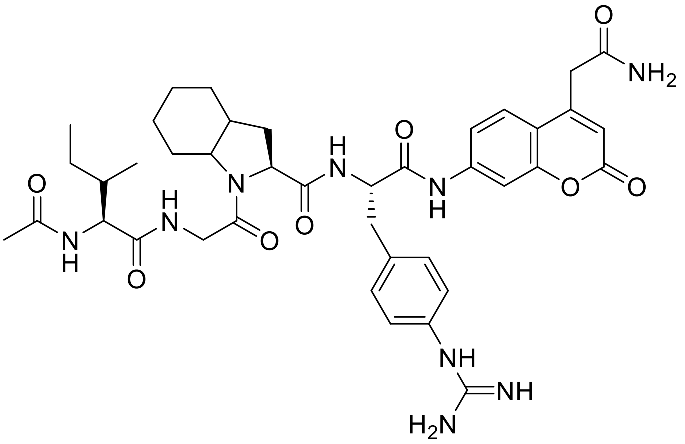


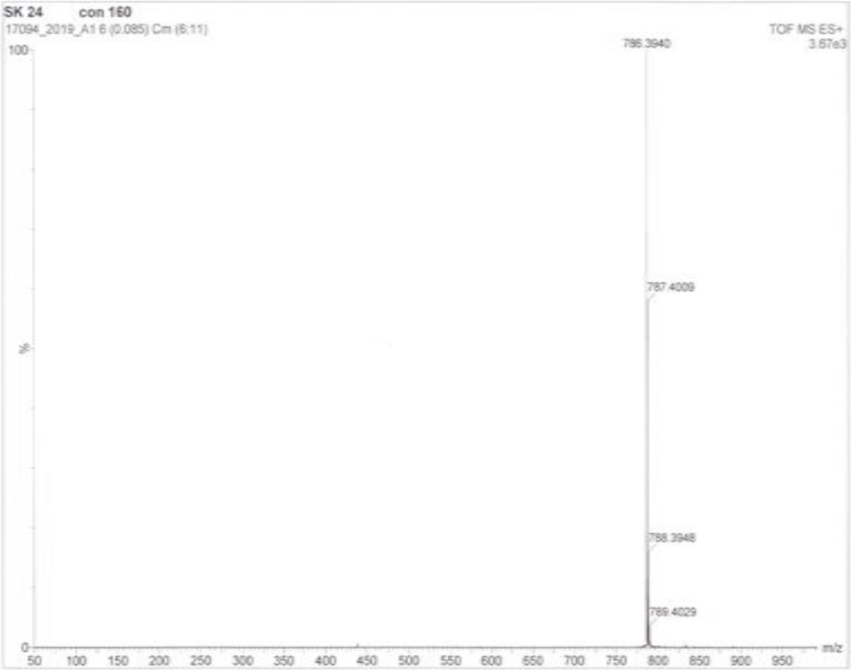


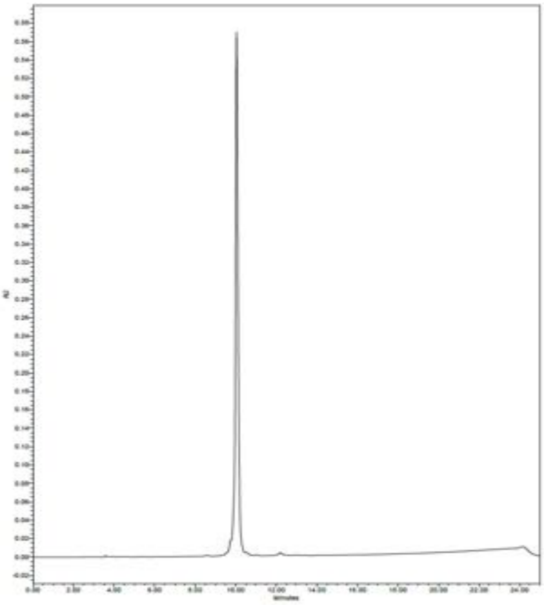

**SK26, Ac-Met(O)-Gly-Tyr(2,6-$Cl_2$-Bzl)-Phe(guan)-ACC**

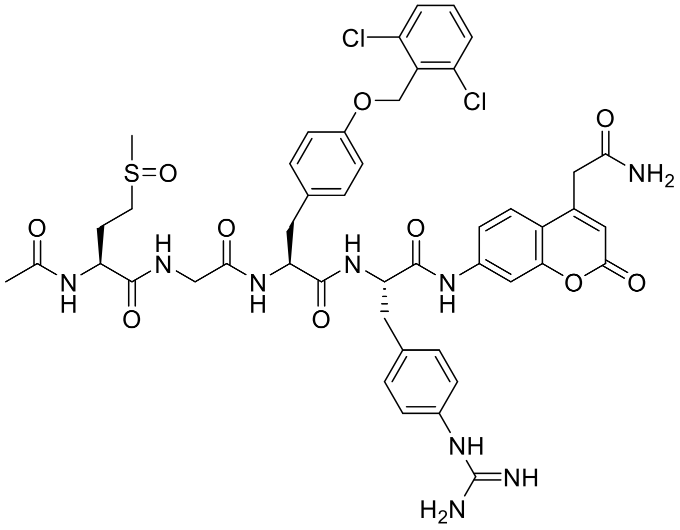

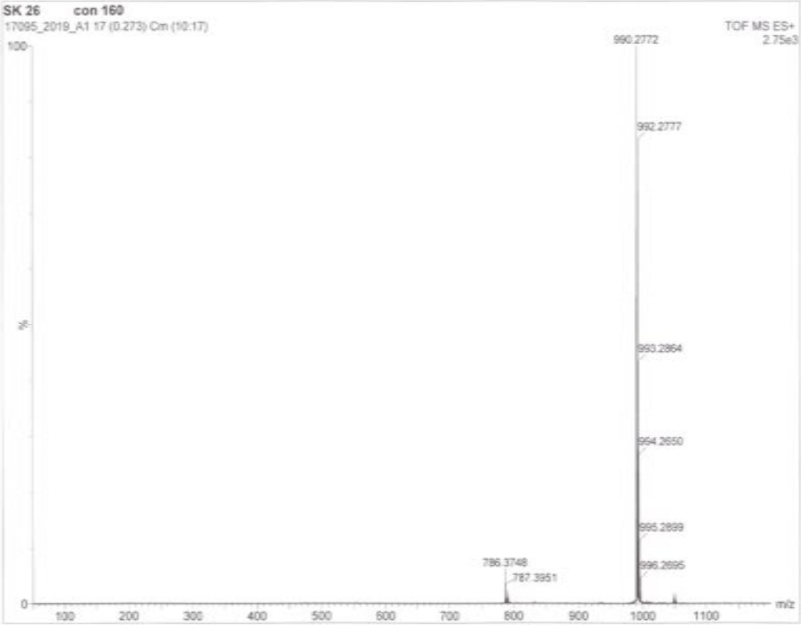


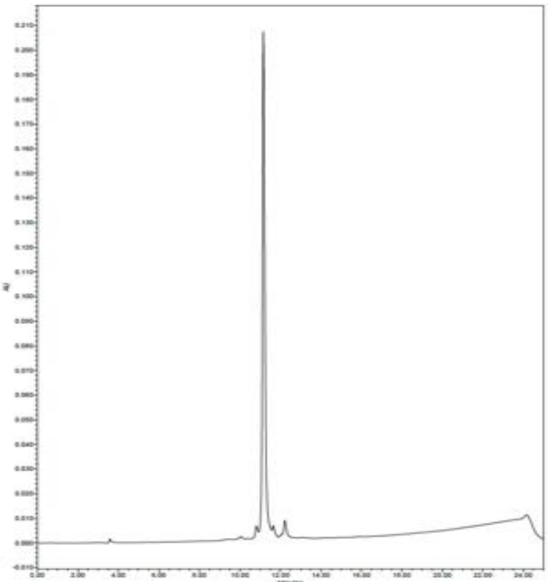

**SK27, Ac-Met(O)-Gly-Hyp(Bzl)-Phe(guan)-ACC**

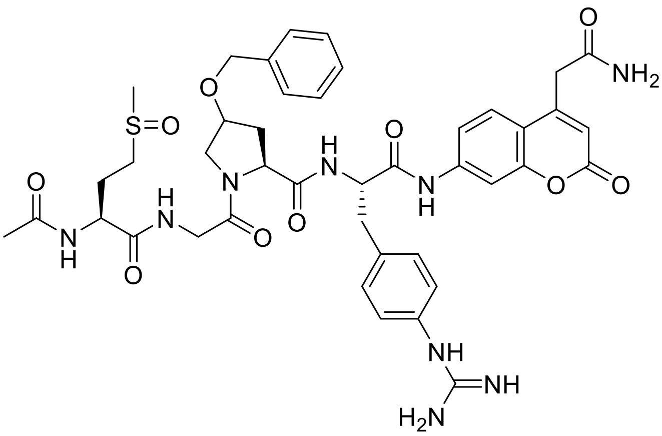

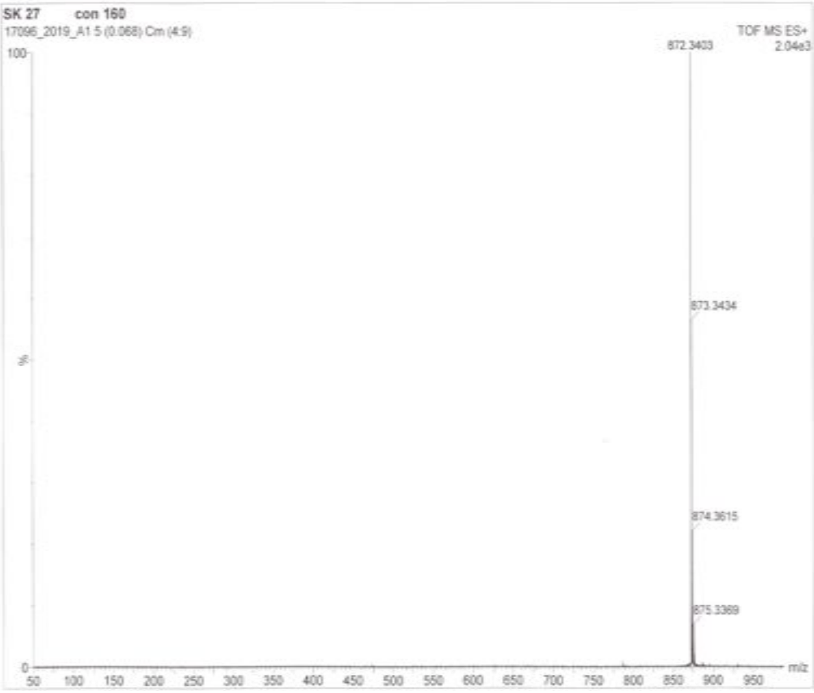


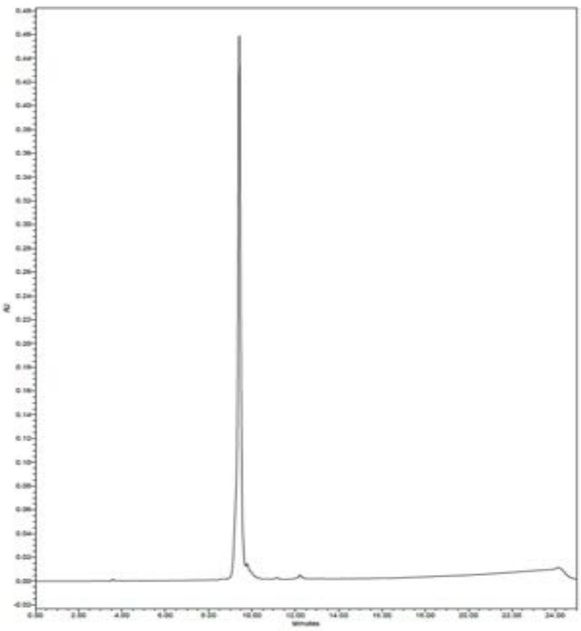

**SK28, Ac-Met(O)-Gly-Phe(4-F)-Phe(guan)-ACC**

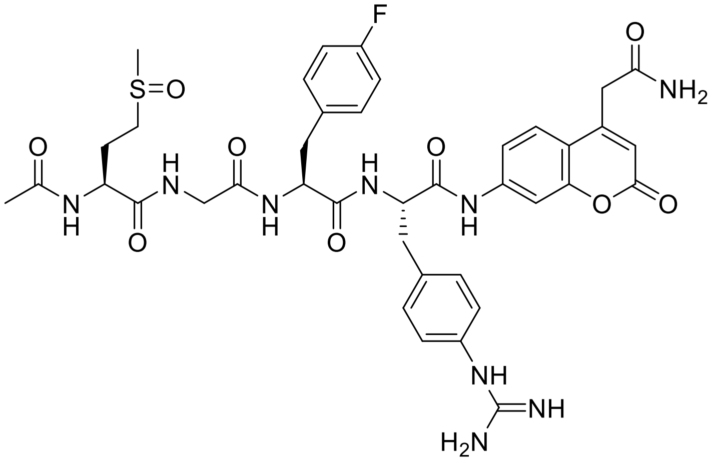


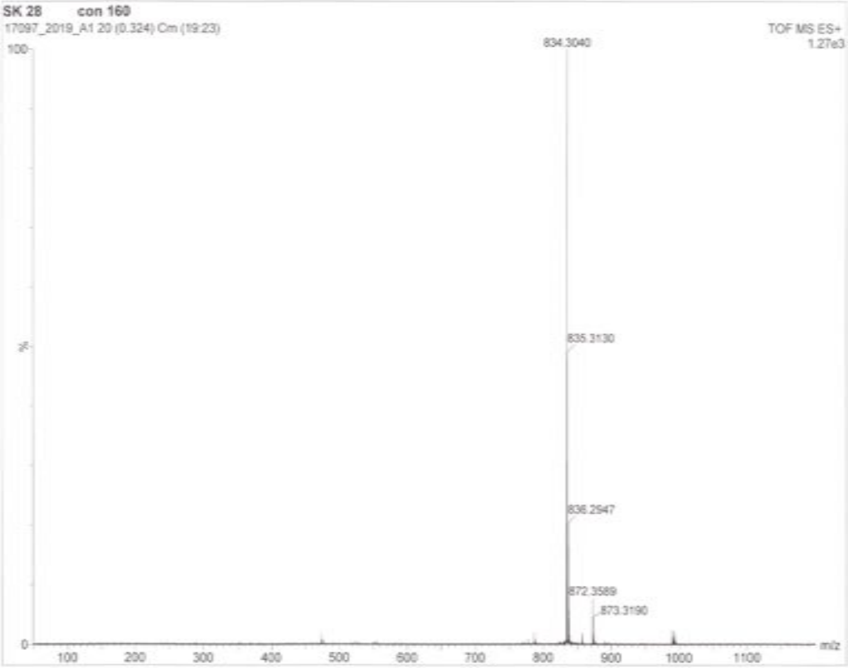


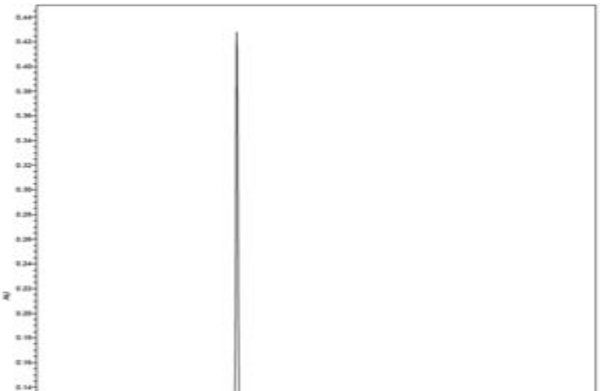

**SK7, Ac-Ile-Gly-Pro-Arg-ACC**

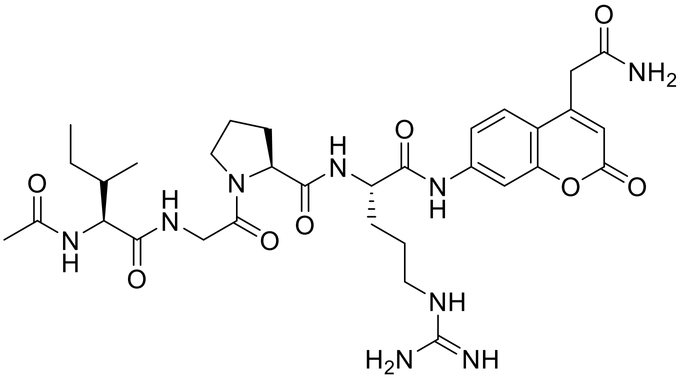

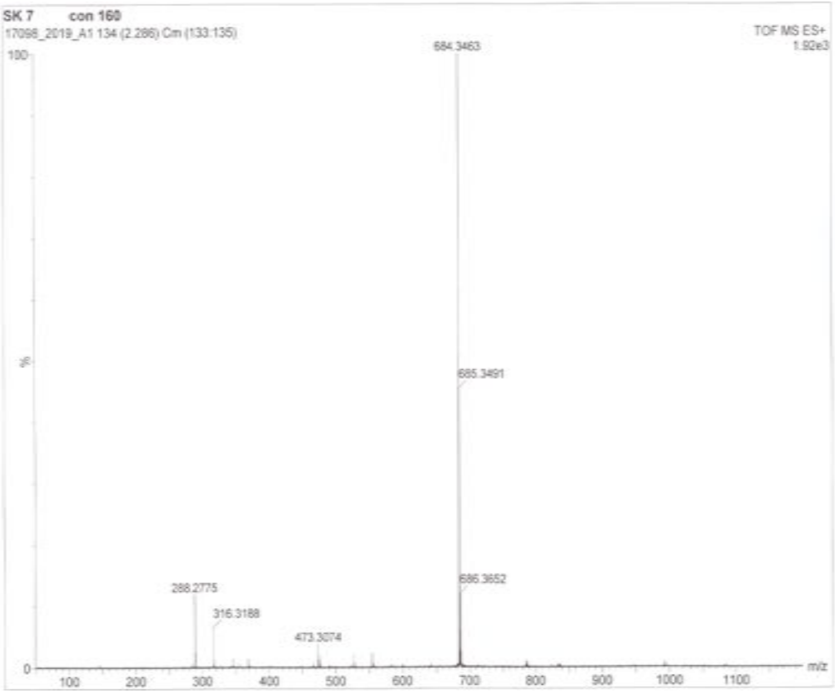


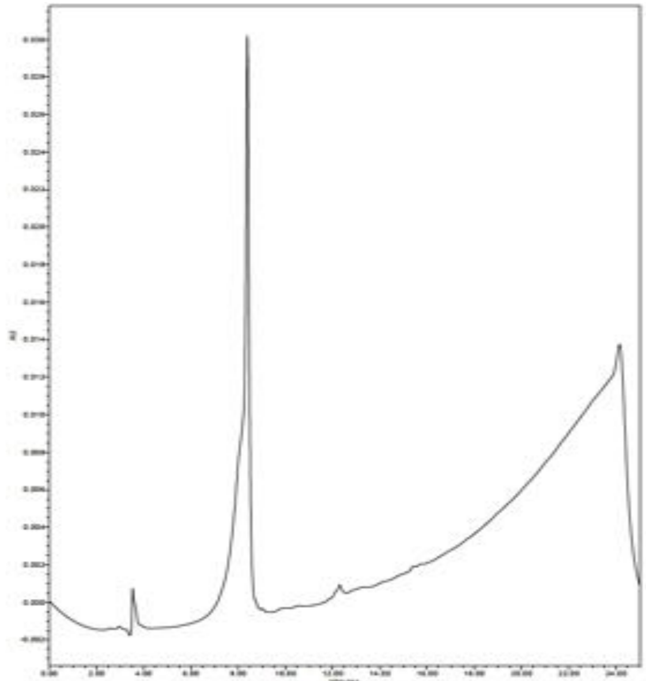

**SK15l, MeOSuc-Tic-Gly-Oic-Arg$^{P}$(OPh)$_2$**

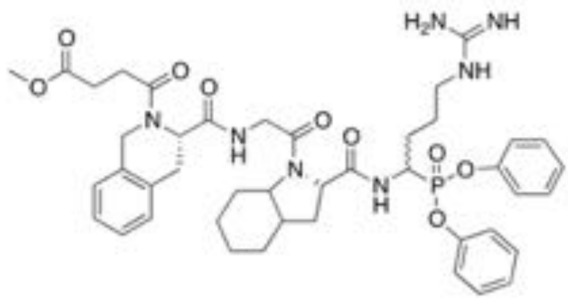


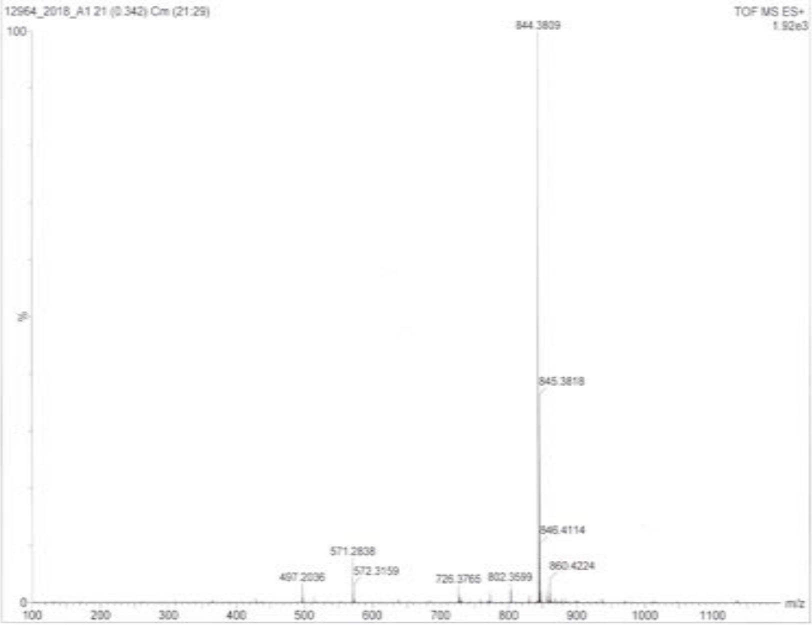


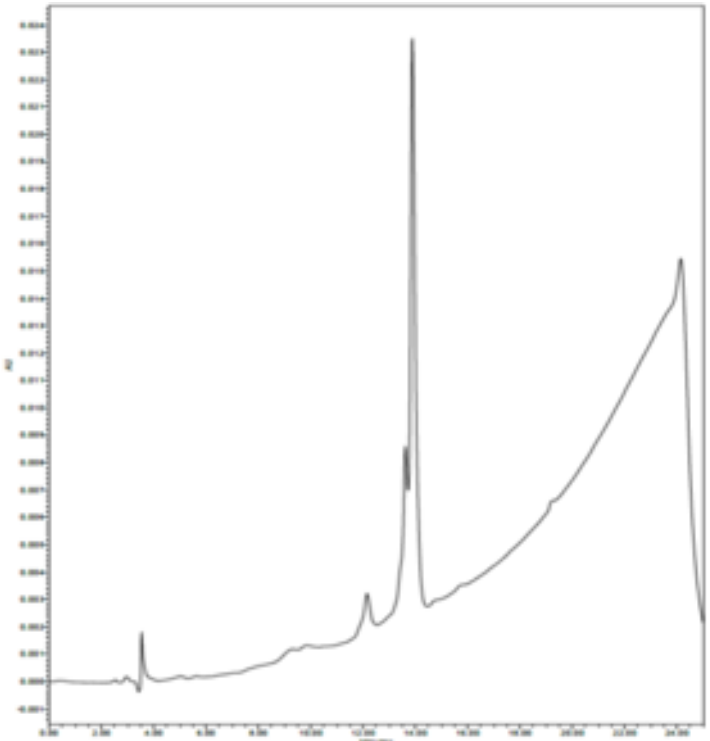

**SK15.5, Cy5-Ahx-Tic-Gly-Oic-Arg$^{P}$(OPh)$_2$**

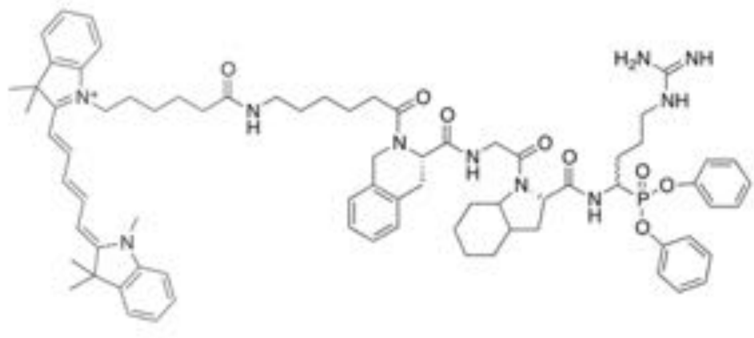

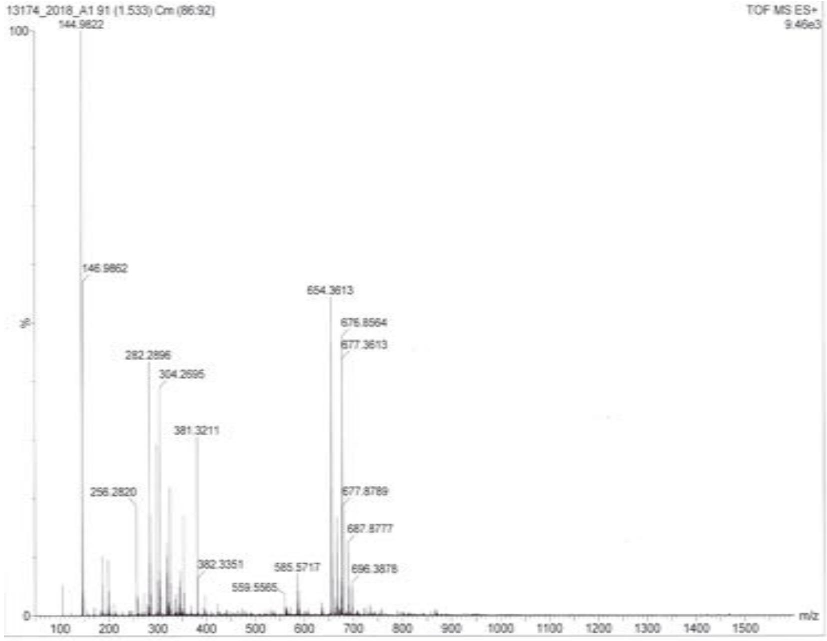


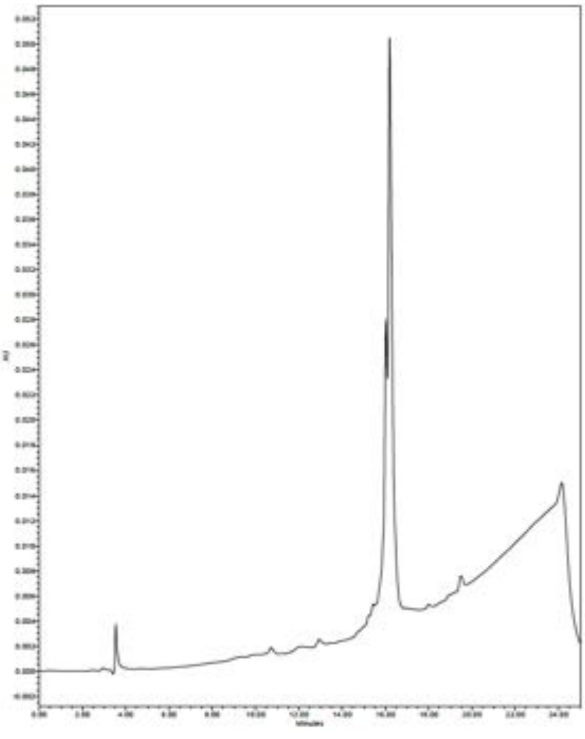

**Figure S1. GrA substrate specificity profiles presented as heat maps.**

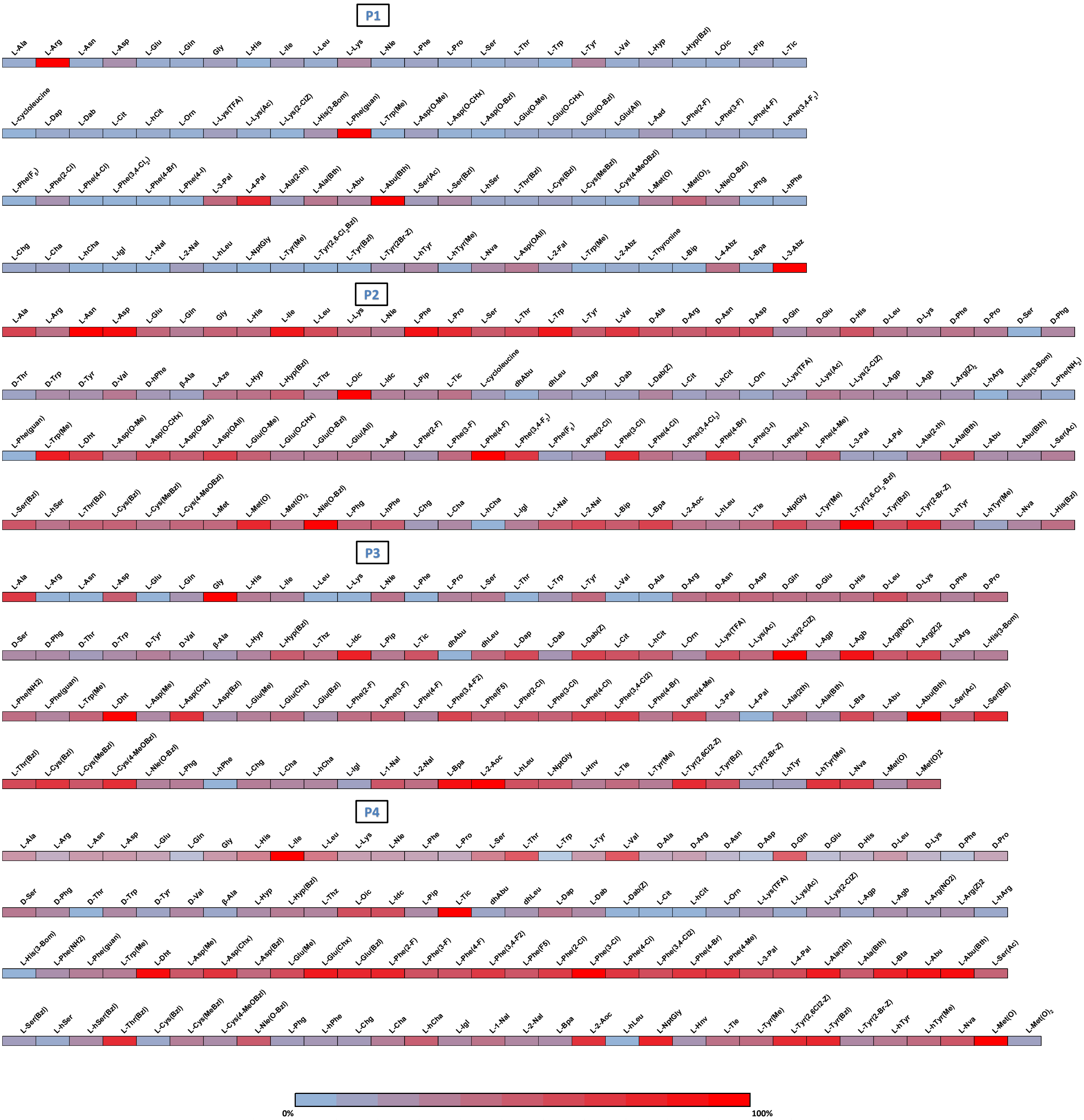

**Table S2. Structures of natural and unnatural amino acids used in P1-Arg HyCoSuL library and Ac-ISPX-ACC library.**

| Lp | Structure + code | Lp | Structure + code | Lp | Structure + code |
|---|---|---|---|---|---|
| 1 | 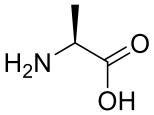 L-Ala | 2 | 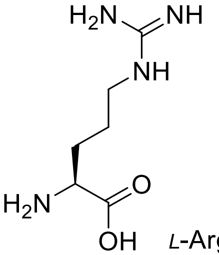 L-Arg | 3 | 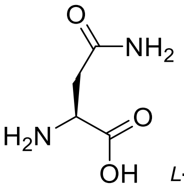 L-Asn |
| 4 | 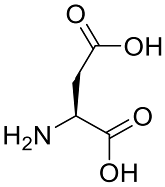 L-Asp | 5 | 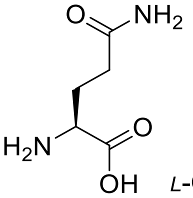 L-Gln | 6 | 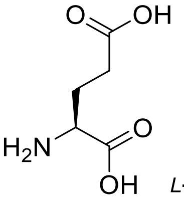 L-Glu |
| 7 | 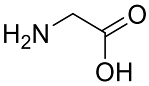 Gly | 8 | 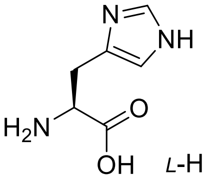 L-His | 9 | 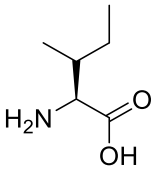 L-Ile |
| 10 | 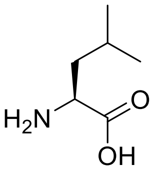 L-Leu | 11 | 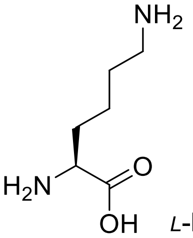 L-Lys | 12 | 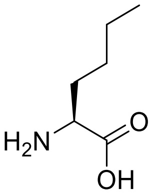 L-Nle |
| 13 | 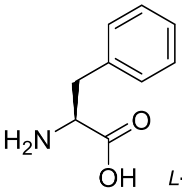 L-Phe | 14 | 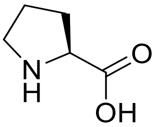 L-Pro | 15 | 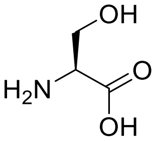 L-Ser |
| 16 | 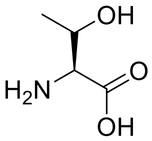 L-Thr | 17 | 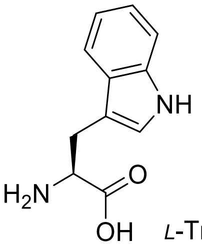 L-Trp | 18 | 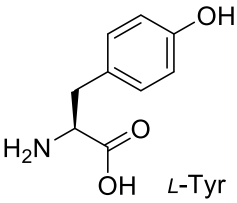 L-Tyr |

19 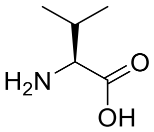 *L*-Val

20 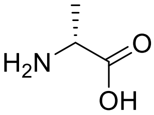 *D*-Ala

21 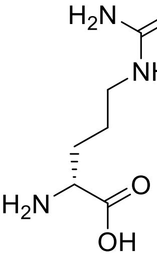 *D*-Arg

22 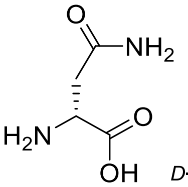 *D*-Asn

23 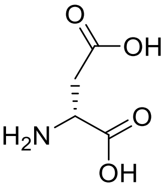 *D*-Asp

24 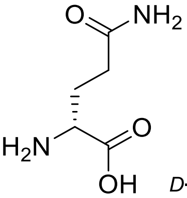 *D*-Gln

25 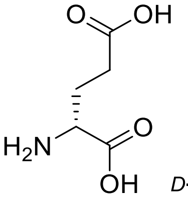 *D*-Glu

26 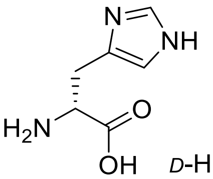 *D*-His

27 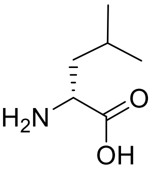 *D*-Leu

28 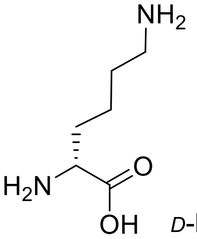 *D*-Lys

29 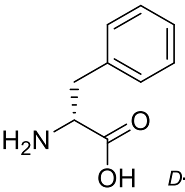 *D*-Phe

30 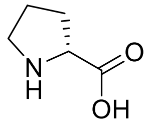 *D*-Pro

31 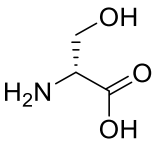 *D*-Ser

32 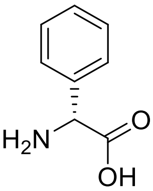 *D*-Phg

33 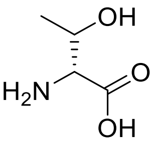 *D*-Thr

34 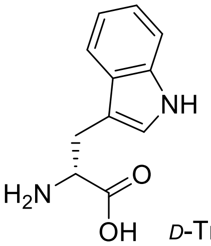 *D*-Trp

35 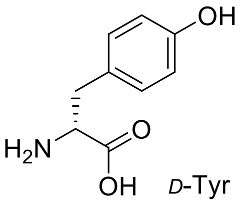 *D*-Tyr

36 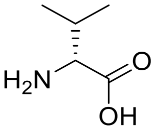 *D*-Val

37 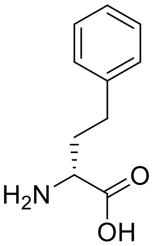 *D*-hPhe

38 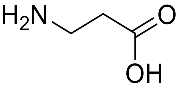 [illegible]-Ala

39 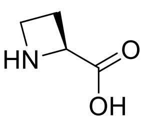 *L*-Aze

40 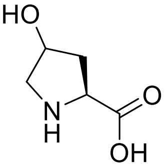 *L*-Hyp

41 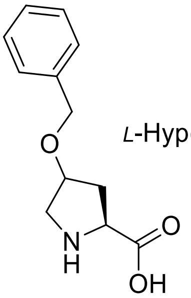 *L*-Hyp(Bzl)

42 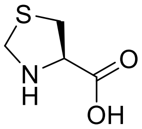 *L*-Thz

43 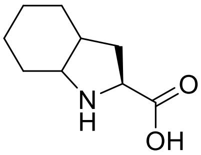 *L*-Oic

44 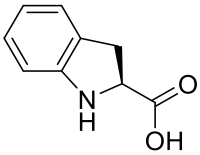 *L*-Idc

45 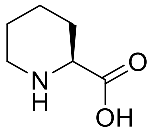 *L*-Pip

46 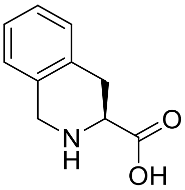 *L*-Tic

47 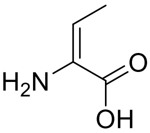 dhAbu

48 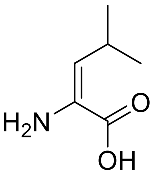 dhLeu

49 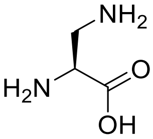 *L*-Dap

50 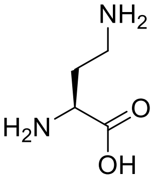 *L*-Dab

51 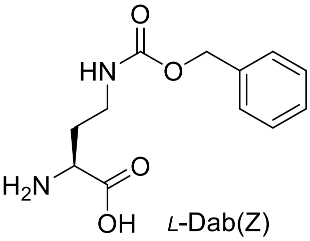 *L*-Dab(Z)

52 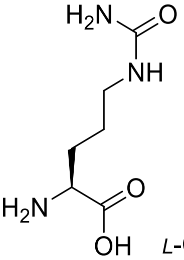 *L*-Cit

53 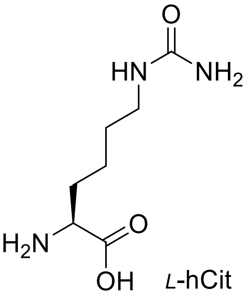 *L*-hCit

54 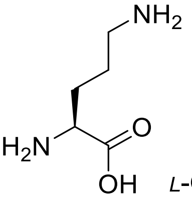 *L*-Orn

55 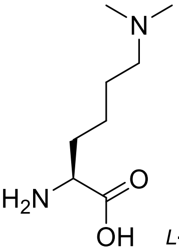 *L*-Lys(Me)$_2$

56 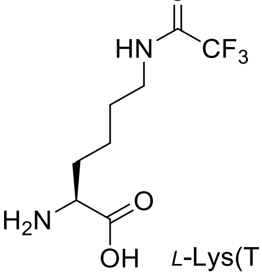 *L*-Lys(TFA)

57 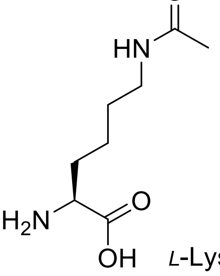 *L*-Lys(Ac)

58 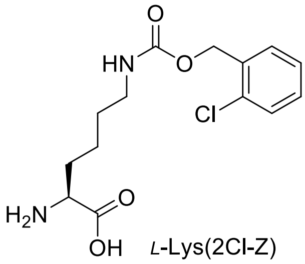 L-Lys(2Cl-Z)

59 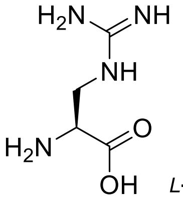 L-Agp

60 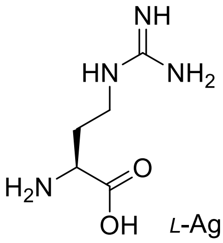 L-Agb

61 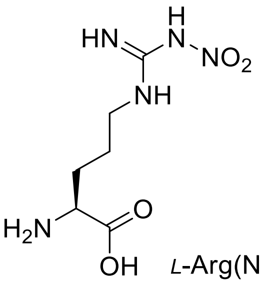 L-Arg($NO_2$)

62 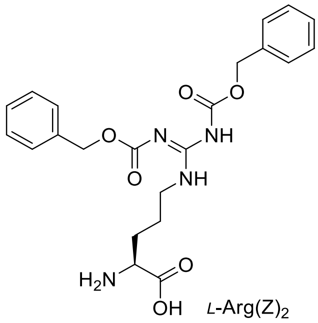 L-Arg$(Z)_2$

63 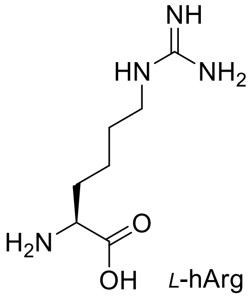 L-hArg

64 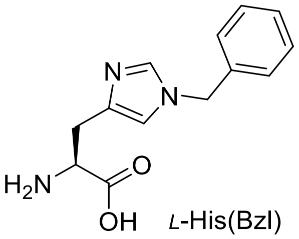 L-His(Bzl)

65 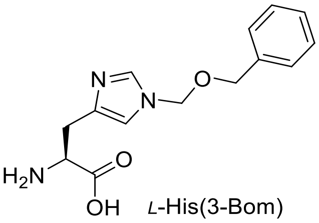 L-His(3-Bom)

66 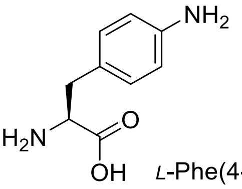 L-Phe(4-$NH_2$)

67 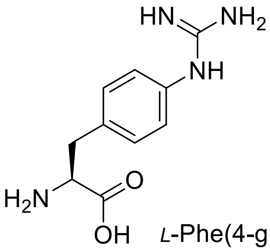 L-Phe(4-guan)

68 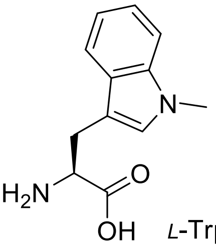 L-Trp(Me)

69 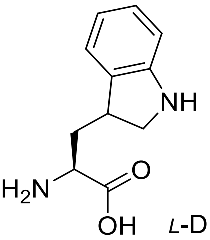 L-Dht

70 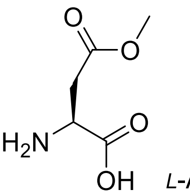 L-Asp(Me)

71 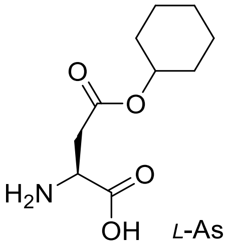 L-Asp(Chx)

72 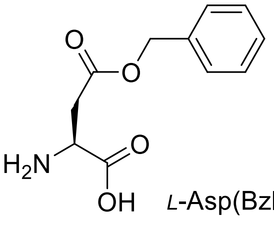 L-Asp(Bzl)

73 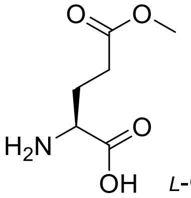 L-Glu(Bzl)

74 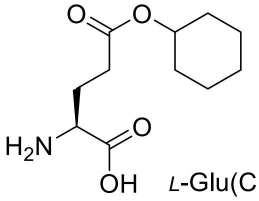 L-Glu(Chx)

75 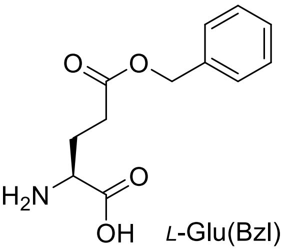 L-Glu(Bzl)

76 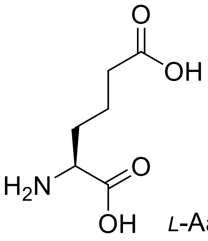 *L*-Aad

77 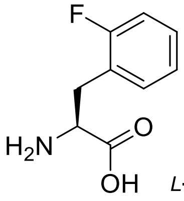 *L*-Phe(2-F)

78 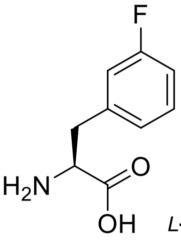 *L*-Phe(3-F)

79 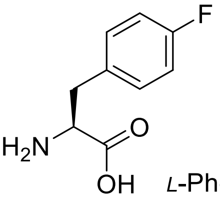 *L*-Phe(4-F)

80 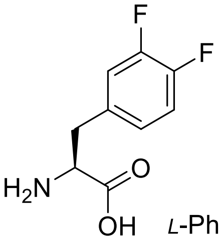 *L*-Phe(4,3-$F_2$)

81 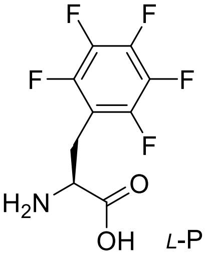 *L*-Phe($F_5$)

82 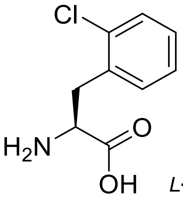 *L*-Phe(2-Cl)

83 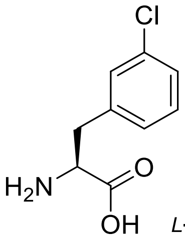 *L*-Phe(3-Cl)

84 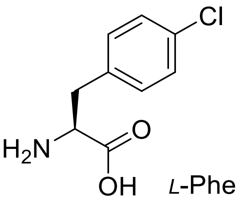 *L*-Phe(4-Cl)

85 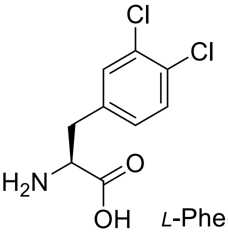 *L*-Phe(3,4-$Cl_2$)

86 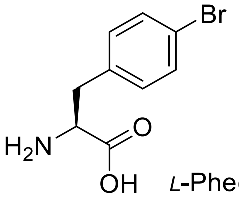 *L*-Phe(4-Br)

87 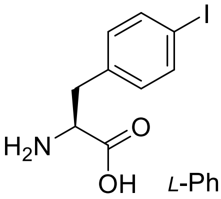 *L*-Phe(4-I)

88 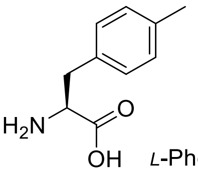 *L*-Phe(4-Me)

89 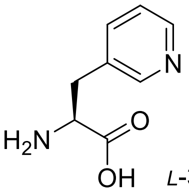 *L*-3-Pal

90 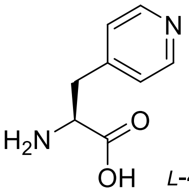 *L*-4-Pal

91 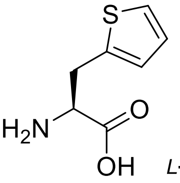 *L*-Ala(2-th)

92 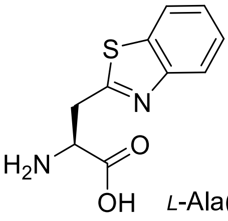 *L*-Ala(Bth)

93 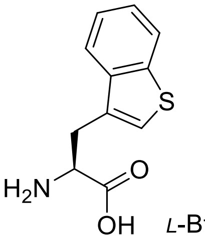 *L*-Bta

94 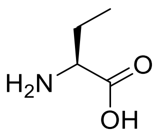 *L*-Abu

95 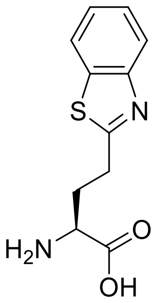 *L*-Abu(Bth)

96 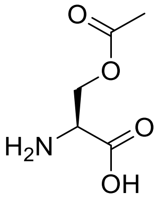 *L*-Ser(Ac)

97 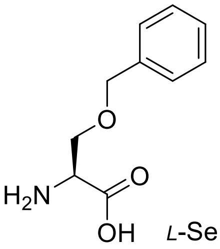 *L*-Ser(Bzl)

98 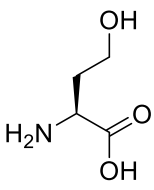 *L*-hSer

99 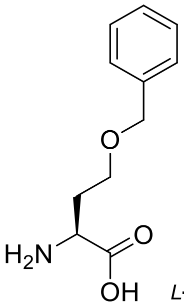 *L*-hSer(Bzl)

100 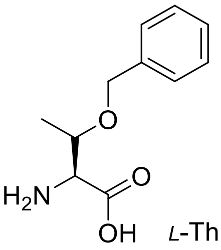 *L*-Thr(Bzl)

101 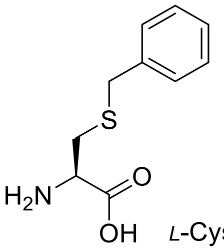 *L*-Cys(Bzl)

102 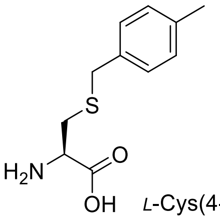 *L*-Cys(4-MeBzl)

103 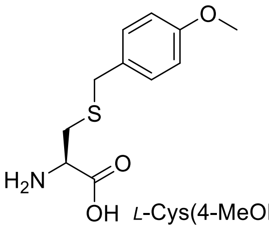 *L*-Cys(4-MeOBzl)

104 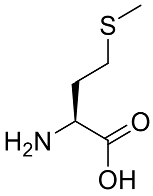 *L*-Met

105 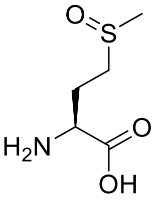 *L*-Met(O)

106 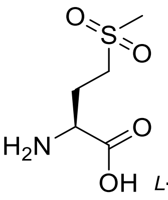 *L*-Met($O_2$)

107 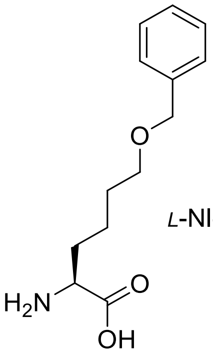 *L*-Nle(*O*-Bzl)

108 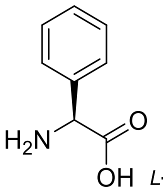 *L*-Phg

109 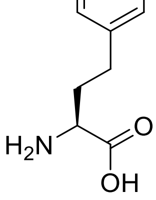 *L*-hPhe

110 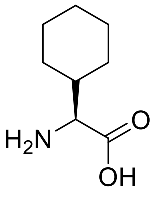 *L*-Chg

111 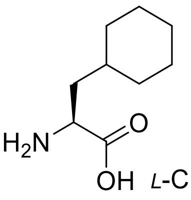 *L*-Cha

112 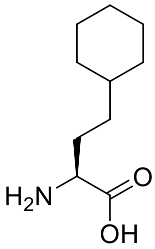 *L*-hCha

113 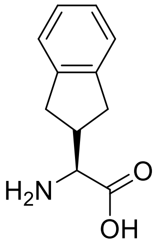 *L*-Igl

114 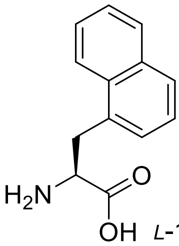 *L*-1-Nal

115 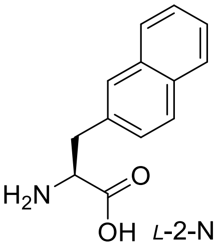 *L*-2-Nal

116 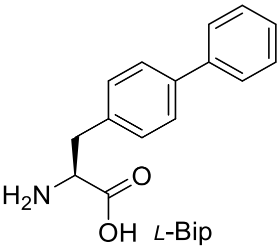 *L*-Bip

117 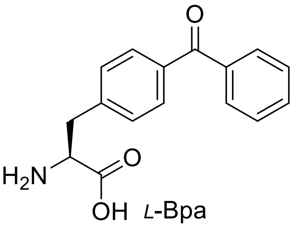 *L*-Bpa

118 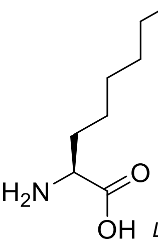 *L*-2-Aoc

119 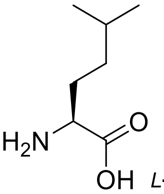 *L*-hLeu

120 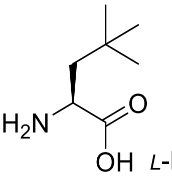 *L*-NptGly

121 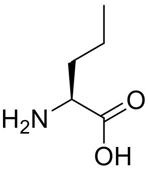 *L*-Nva

122 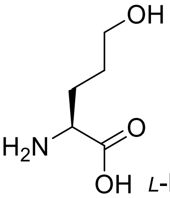 *L*-Hnv

123 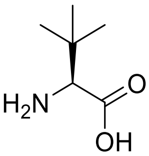 *L*-Tle

124 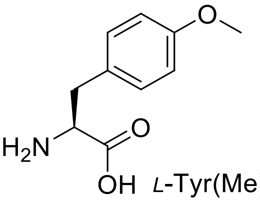 *L*-Tyr(Me)

125 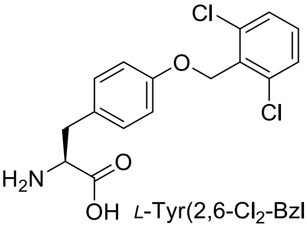 *L*-Tyr(2,6-$Cl_2$-Bzl)

126 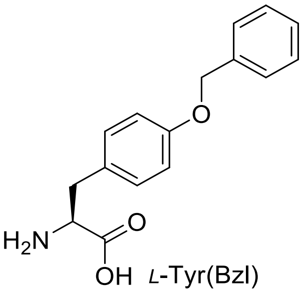 *L*-Tyr(Bzl)

127 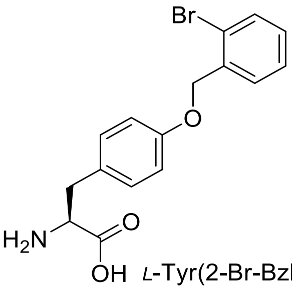 *L*-Tyr(2-Br-Bzl)

128 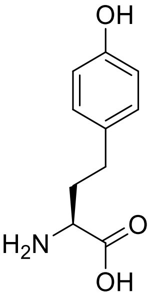 *L*-hTyr

129 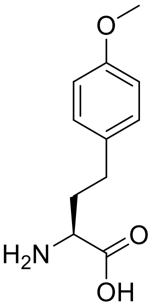 *L*-hTyr(Me)

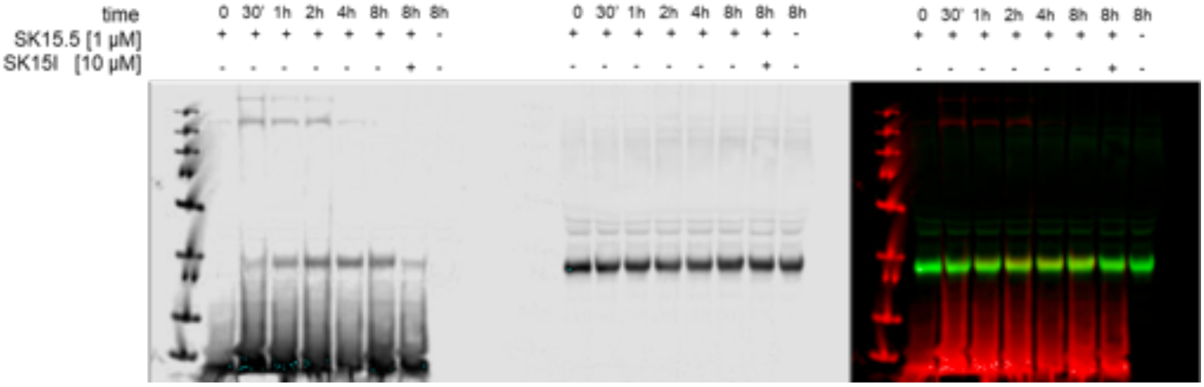


**Figure S2. Detection of active GrA in NK92 cells.**

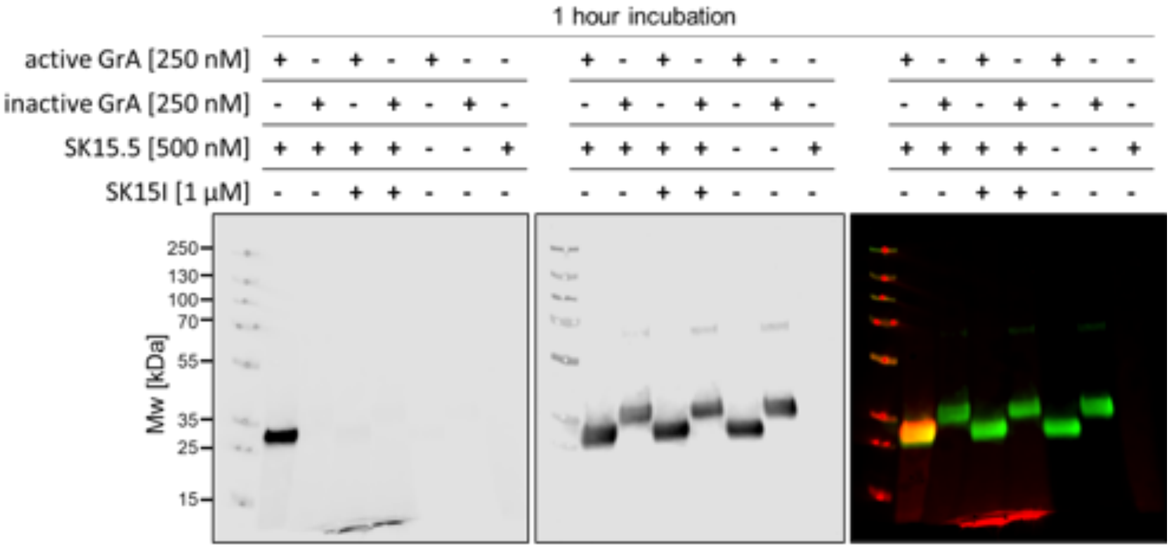


**Figure S3. SK15.5 binds exclusively with the active GrA.**
